# Single molecule array measures of LRRK2 kinase activity in serum link Parkinson’s disease severity to peripheral inflammation

**DOI:** 10.1101/2024.04.15.589570

**Authors:** Yuan Yuan, Huizhong Li, Kashyap Sreeram, Tuyana Malankhanova, Ravindra Boddu, Samuel Strader, Allison Chang, Nicole Bryant, Talene A. Yacoubian, David G. Standaert, Madalynn Erb, Darren J. Moore, Laurie H. Sanders, Michael W. Lutz, Dmitry Velmeshev, Andrew B. West

## Abstract

**Background:** LRRK2-targeting therapeutics that inhibit LRRK2 kinase activity have advanced to clinical trials in idiopathic Parkinson’s disease (iPD). LRRK2 phosphorylates Rab10 on endolysosomes in phagocytic cells to promote some types of immunological responses. The identification of factors that regulate LRRK2-mediated Rab10 phosphorylation in iPD, and whether phosphorylated-Rab10 levels change in different disease states, or with disease progression, may provide insights into the role of Rab10 phosphorylation in iPD and help guide therapeutic strategies targeting this pathway.

**Methods:** Capitalizing on past work demonstrating LRRK2 and phosphorylated-Rab10 interact on vesicles that can shed into biofluids, we developed and validated a high-throughput single-molecule array assay to measure extracellular pT73-Rab10. Ratios of pT73-Rab10 to total Rab10 measured in biobanked serum samples were compared between informative groups of transgenic mice, rats, and a deeply phenotyped cohort of iPD cases and controls. Multivariable and weighted correlation network analyses were used to identify genetic, transcriptomic, clinical, and demographic variables that predict the extracellular pT73-Rab10 to total Rab10 ratio.

**Results:** pT73-Rab10 is absent in serum from *Lrrk2* knockout mice but elevated by *LRRK2* and *VPS35* mutations, as well as *SNCA* expression. Bone-marrow transplantation experiments in mice show that serum pT73-Rab10 levels derive primarily from circulating immune cells. The extracellular ratio of pT73-Rab10 to total Rab10 is dynamic, increasing with inflammation and rapidly decreasing with LRRK2 kinase inhibition. The ratio of pT73-Rab10 to total Rab10 is elevated in iPD patients with greater motor dysfunction, irrespective of disease duration, age, sex, or the usage of PD-related or anti-inflammatory medications. pT73-Rab10 to total Rab10 ratios are associated with neutrophil activation, antigenic responses, and the suppression of platelet activation.

**Conclusions:** The extracellular ratio of pT73-Rab10 to total Rab10 in serum is a novel pharmacodynamic biomarker for LRRK2-linked innate immune activation associated with disease severity in iPD. We propose that those iPD patients with higher serum pT73-Rab10 levels may benefit from LRRK2-targeting therapeutics to mitigate associated deleterious immunological responses.

## Background

Genetic studies implicate missense mutations and promoter variants in the *leucine-rich repeat kinase 2* (*LRRK2*) gene in Parkinson’s disease (PD) risk. Clinical phenotypes associated with *LRRK2* mutations largely overlap with *LRRK2-*negative idiopathic PD (iPD)[1], though carriers of the common G2019S mutation may have a slower motor decline compared to many iPD patients[2]. In genome-wide association studies, the rs76904798 variant in the *LRRK2* promoter increases iPD risk, independent of *LRRK2* missense mutations, while the *LRRK2* N551K/R1398H allele may decrease iPD risk[3,4]. The role of LRRK2 in disease progression in iPD is not clear[5]. Biomarkers for LRRK2 activity, as they may reflect different disease states or change with disease progression, could provide insights into the role of LRRK2 in disease progression. The *LRRK2* gene encodes LRRK2, a serine/threonine kinase with a tandem Rab-like ROC domain and several known substrates. However, there have been very few studies evaluating LRRK2-dependent activity markers in larger cohorts of iPD cases and controls. Since LRRK2-targeting drugs have advanced to iPD, a better understanding of which iPD patients to target, and when, may be critical for successful therapeutic development.

LRRK2 phosphorylates several Rab small GTPase substrates that function in the endolysosomal system in different types of cells throughout the body[6]. The LRRK2 substrate Rab10 is highly expressed in innate immune cells[7]. Activated Rab10 is associated with endolysosomes and has been implicated in endocytosis and vesicle recycling[8,9]. Both LRRK2 and Rab10 have been implicated in mycobacteria infection, TLR4 trafficking, and the regulation of ER dynamics and morphology[10,11]. LRRK2-mediated phosphorylation of Rab10 at threonine 73 (i.e., pT73-Rab10) may promote immune signaling pathways, including PI3K-AKT signaling in cell survival and proliferation, by blocking fast-recycling of Rab10-positive vesicles[7]. Of possible relevance to iPD, aggregated α-synuclein may stimulate the production of pT73-Rab10-positive vesicles in microglia, monocytes and macrophages[12]. Though the common pathogenic *LRRK2* mutation G2019S increases autophosphorylated LRRK2 protein in both mouse models and in vesicle-fractions collected from patient biofluids[13–15], the effects of G2019S-LRRK2 on pT73-Rab10 levels have not been clear, with several studies reporting no effects of the mutation[16,17].

While the role of Rab10 phosphorylation in inflammation phenotypes in iPD is not clear, recent reports suggest that LRRK2 kinase activity in circulating immune cells in the periphery may modify neurodegeneration phenotypes in the brain, though the specific mechanisms underlying these observations have not been elucidated[12,18]. A proportion of LRRK2 and Rab10 proteins are bound to vesicles including those subject to export into the extracellular space[19,20]. Our recent results identified higher pT73-Rab10 levels in extracellular vesicle (EV) fractions purified from urine in iPD cases with worse disease phenotypes[21]. However, pT73-Rab10 and total Rab10 proteins were measured in that study by western blots normalized to baseline values, as absolute ratios of pT73-Rab10 to total Rab10 could not be determined.

Further, past biomarker approaches for LRRK2 or LRRK2-phosphorylated Rab proteins have required large volumes of urine (e.g., >40 mL per subject) or freshly-donated blood to isolate and select lysates from certain cell populations or enriched EV fractions. To overcome some of the past challenges associated with high-throughput measures of LRRK2 activity and Rab phosphorylation in patient samples, here we develop an ultra-sensitive version of a SiMOA-based approach, focusing on the analysis of pT73-Rab10 in serum. With specificity and sensitivity defined in-part through the analysis of serum from well-characterized transgenic mouse and rat strains, our results in both rodents and human samples establish new links between extracellular LRRK2 levels, pT73-Rab10, and antigenic myeloid cell responses that correlate with worse motor defects in idiopathic PD.

## Methods

### SiMOA assays

All reagents for the described single-molecule assays used in this study are commercially available. For up-to-date protocols for the measurement of total LRRK2, total Rab10, and pT73-Rab10, detailed graphical step-by-step procedures are hosted on Protocols.Io and available through the following links:

dx.doi.org/10.17504/protocols.io.dm6gp3188vzp/v1 (total Rab10)
dx.doi.org/10.17504/protocols.io.e6nvwdn69lmk/v1 (pT73-Rab10)
dx.doi.org/10.17504/protocols.io.yxmvm317ol3p/v1 (total LRRK2)

Antibodies used for protein capture include anti-LRRK2 (N241A/34, Antibodies Inc. USA), anti-panRab10 (clone 605B11, NanoTools, Germany) and anti-pT73 Rab10 (clone MJF-R2, ab231707, Abcam, USA). Detection antibodies include anti-LRRK2 [MJFF2 (c41-2)] (ab172378, Abcam, USA) and anti-RAB10 [MJF-R23] (ab237703, Abcam, USA). Antibody stock concentration was determinted with a Nanodrop one spectrophotometer (ThermoFisher Scientific, USA). Initial recombinant protein calibrators used in this study that can also be used as protein standards spiked into relevant detection matrices included LRRK2 [G2019S] recombinant human protein (ab177261, Abcam) and recombinant Rab10 (NBP2-23392, Novus Biologicals). All sample incubations, washing, and detection steps are performed on a Quanterix SR-X system and all results analyzed as AEB (average enzymes per bead) values fitted to an optimized non-linear regression curve for each run.

### Cell culture and western blots

HEK293T cells grown in DMEM (Gibco, USA) supplemented with 10% fetal bovine serum (Gibco, USA) were transfected with pcDNA3.1-FLAG-LRRK2 and Rab10 plasmids with polyethylenimine hydrochloride (PEI, Polyscience, USA) and lysates harvested ∼16-hours later. As indicated, cells were treated with 200 nM MLi2 (Sigma, USA) for 2 hours prior to lysis. Cells were lysed in 50mM Tris-HCl pH 8.0 buffer with 150mM NaCl, 1% Triton 100x, and Complete, Protease Inhibitor and PhosSTOP phosphatase inhibitor (Roche, USA). Cell lysates centrifuged at 10 xk*g* for 10 min at 4°C, and protein concentration in supernatant determined by BCA assay (Thermo Fisher Scientific, USA).

Mouse bone marrow-derived macrophages (BMDMs) were collected from the femur and tibia from 3- to 5-month-old mice and grown in DMEM (Gibco, USA) supplemented with 10% fetal bovine serum (Gibco, USA) and 100 ng per mL M-CSF (BioLegend, USA). For western blot analysis, cells were lysed directly in 2x Laemmli sample buffer (4% SDS, 20% glycerol, 0.004% bromophenol blue, 0.125M Tris-HCL-pH6.8, 10% DTT) supplemented with Protease and PhosSTOP inhibitor tablets (Roche, USA).

Samples were electrophoresed on BioRad mini-PROTEAN TGX Stain-Free 4-20%,15-well gradient gels (BioRad, USA), and transferred toPVDF membranes (Sigma, USA). Phos-tag gels were formulated using the Phos-Tag Common Solution (FUJIFILM Wako, USA) as recommended by the manufacturer. Separating gels were used with 12% acrylamide and 5 mM Phos-tag reagent along with 1.5M Tris-HCl, pH=8.8,10mM MnCl_2_, 10% SDS, 10% ammonium persulfate (APS), and 1% tetramethylethylenediamine (TEMED). Separating gels were cast on BioRad mini-PROTEAN short plates (BioRad, USA). Stacking gels were formulated with 10% acrylamide, 0.5M Tris-HCl, pH=6.8, 10% SDS, 10% APS, and 1% TEMED. Gels were electrophoresed at constant 100V for approximately 2 hrs, soaked in a running buffer supplemented with 1 mM EDTA, and then transferred to PVDF. Primary antibodies utilized include pT73-Rab10 (ab231707, abcam), total-Rab10 (ab237703, abcam), and total-LRRK2 (N241 75-253, NanoTools).

### Procedures in rodents

Animal protocols were approved by local Animal Care and Use Committees accredited by the AAALAC. Tg-WT-*Lrrk2* (JAX stock #012466), Tg-G2019S BAC-*Lrrk2* (JAX stock #012467), *Lrrk2*^R1441C/R1441C^ (JAX stock #009346), *VPS35*^WT/WT^, *VPS35*^WT/D620N^ and *VPS35*^D620N/D620N^ (JAX stock #023409), *Lrrk2^-/-^* knock-out (JAX stock #016121), C57BL/6J (JAX stock #000664), BL/6-CD45.1 (JAX stock#002014), Snca^-/-^ (JAX stock#016123) and PAC-SNCA/Snca^-/-^(JAX stock#010710) strains were obtained from Jackson Laboratories. CD1 (CR #022CD1) mice and outbred Long Evans *Lrrk2^+/+^* rats were obtained from Charles River Laboratories. *Lrrk2* knock-out Long-Evans rats were obtained from Envigo. Venous serum was collected in mice via facial vein draws, and in rats via lateral saphenous vein draws, each into microtainer serum collection tubes (Becton Dickinson, USA). 30 min post-collection, tubes were centrifuged at 8 xk*g* for 5 min at RT and serum supernatants aliquoted and stored at -80°C. Serum samples with less than 40 µL available, or showing evidence of extensive hemolysis of more than 100 mg per dL hemoglobin, as estimated by standard scales, were excluded from further analysis.

The small molecule LRRK2 kinase inhibitor PFE-360 (PF-06685360, Pharmaron, USA) was administered to rodents in a suspension solution consisting of deionized water and 0.5% (w/v) methylcellulose (Sigma, USA). Drug solutions were made the day before administration with 4°C incubation overnight with stir-bar mixing. Formulations were administered with 18-gauge curved 2.4 mm ball point stainless feeding needles (KVP International, USA). Rodents were dosed at 10 mg per kg. The drug suspension was prepared at 1 mg per mL for mice and 5 mg per mL for rats.

For whole body irradiation, mice were exposed to 11 grays (split into two equal doses separated by 3 hrs) using a Xstrahl CIX3 Xray irradiator and maintained on acidified-antibiotic water consisting of NaOCl, HCl, and neomycin sulfate for 6 days. For bone marrow transplantation, donor mouse cells were isolated from femur and tibia washes with a 26 gauge needle and RPMI1640 media (Gibco, USA) and filtered through a 70 µm cell strainer as described [22]. After centrifugation at 300 xg for 10 min at 4°C, cell pellets were treated with Ammonium-chloride–potassium (ACK) lysis buffer (Gibco, USA). Pelleted cells were then resuspended into fresh RPMI1640 media and immediately transplanted (tail vein injection) with 3-5×10^6^ bone marrow mononuclear cells after the second dose of radiation in the host mice. Transplanted mice were maintained on the acidified-antibiotic water for 4 weeks.

Flow cytometry analysis of mouse blood was accomplished with freshly collected blood (via facial vein puncture) with RBC lysis for 10 min at RT in ACK solution, and centrifugation at 300 x*g* for 10 min at 4°C as described [23,24]. Leukocyte pellets were washed and suspended in buffer (PBS, 0.5% bovine serum albumin, and 0.01% sodium azide), blocked with Fcγ receptor antibody (clone 2.4G2, eBioscience, USA), and cells stained with brilliant violet-CD45 (clone 30-F11, Biolegend, USA); allophycocyanin-conjugated-CD45.1 (clone A20, eBioscience, USA); fluorescein isothiocyanate-conjugated-CD45.2 (clone 104, eBioscience, USA); allophycocyanin-conjugated Ly6G (Gr-1, clone 1A8, eBioscience, USA); eFlour 450-conjugated Ly6C (clone HK1.4, eBioscience, USA) and fluorescein isothiocyanate-conjugated CD4 (clone GK1.5, eBioscience, USA). 7-aminoactinomycin D (00-6993-50, Invitrogen, USA) was used for live-dead staining and analyzed on an Attune NxT acoustic focusing instrument (ThermoFisher Scientific, USA) and FlowJo 10.8.1 software (Becton Dickinson, USA).

For polymicrobial sepsis induction, 200 µL of a rapidly thawed cecal slurry preparation was injected intraperitoneal in mice, with serum collections 8 and 72 hours (as indicated) post injection. For the cecal slurry preparation, cecal contents were harvested and pooled from 25 male CD1 mice and mixed at a ratio of 0.5 mL of water to 100 mg of cecal contents, sequentially filtered through a 860 µm,190 µm and 70 µm mesh (Fisher Scientific, USA), and then added to an equal volume of 30% glycerol in PBS as described [25]. Samples were submitted to the University of North Carolina Microbiome Core Facility (supported in part by NIH grant P30 DK34987) for 16S rRNA amplicon sequencing. To determine colony formation unit (CFU) per mL, slurries were plated onto brain-heart-infusion broth agar plates (1.5% agar) and incubated for 24 hours at 37°C in ambient air.

### Human subjects, clinical, and demographic data

This study was approved by local Institutional Review Boards and all participants signed informed consents. All coded serum samples were processed by investigators blinded to sample identity, and final curated datasets were achieved before associating clinical and demographic characteristics were assigned to biomarker values. Serum was obtained from participants (Table 1) all enrolled at a single site at the University of Alabama at Birmingham (UAB) Movement Disorder Clinic, part of the Parkinson’s Disease Biomarker Program (http://pdbp.ninds.nih.gov). Participants were selected based on frequency matching of age and gender between cases and controls, with inclusion for cases based on United Kingdom Parkinson’s Disease Society Brain Bank Criteria. Inclusion criteria for healthy controls were aged 35 to 85 years, lack of PD in first-degree blood relatives, and a lack of positive responses on more than 3 items in the PD Screening Questionnaire[26]. Exclusion criteria for all subjects included atypical features indicative of progressive supranuclear palsy, multiple system atrophy, corticobasal degeneration, cerebellar signs, supranuclear gaze palsy, apraxia, disabling autonomic failure, neuroleptic treatment at time of onset of parkinsonism, active treatment with a neuroleptic at time of study entry, history of repeated strokes with stepwise progression of parkinsonism, history of repeated head injury, history of definite encephalitis, dementia, known severe anemia, or kidney disease as determined by measurement of serum creatinine and/or medical records, infections, or any serious comorbidity that would interfere with participation in the study. Demographics, PD characteristics, medications, medical history, and family history of PD were collected. Levodopa equivalent daily dose was calculated as described[26]. Serum samples were collected according to BioSpecimen Exchange for Neurology Disorders

**Table 1.** Demographics, clinical information, and biomarker data in subjects recruited into the Parkinson’s disease biomarker program (PDBP).

|  | Male (N=276) |  |  | Female (N=246) |  |  |
| --- | --- | --- | --- | --- | --- | --- |
| Cohort feature | PD (N=187) | HC (N=89) | p* | PD (N=90) | HC (N=156) | p* |
| Disease duration, Mean years (SD) | 8.1 (5.9) | 0 | <0.0001 | 6.5 (4.5) | 0 | <0.0001 |
| Age, Mean years (SD) | 64.0 (9.6) | 61.0 (13.2) | 0.259 | 63.9 (8.8) | 59.6 (9.9) | 0.001 |
| MDS-UPDRS I, Mean (SD) | 10.8 (5.9) | 5.1 (4.5) | <0.0001 | 11.8 (6.1) | 5.4 (4.0) | <0.0001 |
| MDS-UPDRS II, Mean (SD) | 14.6 (8.1) | 1.3 (2.6) | <0.0001 | 12.8 (8.3) | 0.9 (2.1) | <0.0001 |
| MDS-UPDRS III, Mean (SD) | 29.1 (13.4) | 0.3 (1.0) | <0.0001 | 24.5 (11.1) | 0.2 (0.7) | <0.0001 |
| MDS-UPDRS IV, Mean (SD) | 1.8 (3.2) | 0 | <0.0001 | 1.9 (2.9) | 0 | <0.0001 |
| MDS-UPDRS Total, Mean (SD) | 56.4 (24.2) | 6.9 (6.8) | <0.0001 | 50.9 (21.1) | 6.4 (5.7) | <0.0001 |
| MoCA, Mean (SD) | 24.0 (3.5) | 25.5 (3.0) | 0.001 | 25.1 (3.0) | 27.1 (1.9) | <0.0001 |
| UPSIT, Mean (SD) | 18.3 (7.3) | 30.9 (6.0) | <0.0001 | 20.4 (8.1) | 34.3 (4.1) | <0.0001 |
| Epworth Sleep Scale, Mean (SD) | 9.9 (4.7) | 6.6 (3.6) | <0.0001 | 9.1 (5.2) | 5.2 (3.2) | <0.0001 |
| NSAID use, N (%) | 84 (45) | 28 (31) | 0.036 | 36 (40) | 46 (29) | 0.095 |
| Statin use, N (%) | 60 (32) | 31 (35) | 0.682 | 25 (28) | 40 (26) | 0.765 |
| LEDD, Mean (SD) | 561 (500) | 0 | <0.0001 | 513 (757) | 0 | <0.0001 |
| Family history of PD, N (%) | 49 (26) | 7 (8) | 0.0003 | 27 (30) | 9 (6) | <0.0001 |
| Serum biomarkers and characteristics |  |  |  |  |  |  |
| Sample days-in-freezer, Mean (SD) | 2214 (364) | 2240 (338) | 0.671 | 2296 (335) | 2184 (344) | 0.007 |
| Total LRRK2, ng/mL, Median (IQR) | 0.77 (2.04) | 0.50 (1.40) | 0.034 | 0.49 (2.43) | 0.50 (1.16) | 0.318 |
| Mean | 12.05 | 4.36 |  | 3.24 | 3.07 |  |
| 95%CI [LL, UL] | [-1.93, 26.02] | [2.08, 6.65] |  | [1.87, 4.63] | [1.33, 4.81] |  |
| pT73-Rab10, ng/mL, Median (IQR) | 1.09 (8.81) | 0.68 (5.77) | 0.252 | 1.19 (9.45) | 0.28 (2.36) | 0.073 |
| Mean | 17.36 | 11.83 |  | 8.69 | 5.70 |  |
| 95%CI [LL, UL] | [7.00, 27.72] | [5.59, 18.07] |  | [5.03, 12.35] | [3.49, 7.91] |  |
| Rab10, ng/mL, Median (IQR) | 7.69 (40.67) | 8.01 (25.46) | 0.491 | 10.51 (47.12) | 5.68 (12.65) | 0.012 |
| Mean | 115.45 | 66.22 |  | 68.24 | 45.77 |  |
| 95%CI [LL, UL] | [60.99, 169.90] | [29.69, 102.75] |  | [38.00, 98.48] | [23.31, 68.23] |  |
| pT73-Rab/Rab10, %, Median (IQR) | 10.1 (23.8) | 8.8 (20.57) | 0.260 | 7.2 (17.8) | 6.2 (14.2) | 0.452 |
| Mean | 17.6 | 15.2 |  | 14.2 | 12.6 |  |
| 95%CI [LL, UL] | [14.5, 20.7] | [10.8, 19.6] |  | [10.3, 18.2] | [9.6, 15.6] |  |

\*p values are from 2-tailed t-tests for all continuous cohort variables and Fisher's exact test for cohort categorical variables. P values from serum biomarkers (non-transformed) are from Mann-Whitney tests.

(BioSEND) protocol. Venous blood was collected in red-capped BD Vacutainer serum tubes that were incubated at RT for 30 min, and then centrifuged for 15 min at 1500 x g at 4°C. Serum samples demonstrating more than 100 mg per dL hemoglobin estimated by visual inspection against colorimetric scales were excluded from analysis. Resultant supernatants were aliquoted and stored at -80°C until further use. Venous blood was next collected into PAXgene blood RNA tubes (IVD, Qiagen, USA) following manufacturer’s recommended protocol. Samples and all clinical data including the MDS-UPDRS were collected between 08:00 and 12:00 in the day with PD patients taking their usual medications. All clinical and demographic data collected from these participants has been deposited into the Data Management Resource managed and available from the National Institutes of Health (DMR, https://pdbp.ninds.nih.gov/portal).

### Serum processing and proteomic analysis

Some serum samples were processed using Exo-Spin serum mini column exosome purification kits (Cell Guidance Systems LLC, USA), where 100 µl of serum samples were centrifuged at 300 x*g* for 10 min at 4°C to remove any large debris and then at 16 xk*g* for 30 min, and supernatant was transferred to the PBS-equilibrated columns. The first flow-through was discarded, and another 180 µL of PBS was added to the top of the column, centrifuged at low speed 20 x*g* for 1 min, fraction 1 collected, and this procedure was repeated two more times. 100 µg of protein (BCA analysis) was processed and evaluated on a Thermo Orbitrap Fusion Lumos mass spectrometer coupled with ultra-performance liquid chromatography (Thermo Fisher, USA). Tandem mass spectra were extracted and all samples were analyzed using Sequest set to search MasterMix_011320.fasta; SP_Hsapiens_082922.fasta. Scaffold (version Scaffold_5.2.2, Proteome Software Inc., USA) was used to validate MS/MS based peptide and protein identifications, where peptide identifications were accepted if they could be established at greater than 92.0% probability to achieve an FDR less than 1.0% by the Percolator posterior error probability calculation [27]. Protein identifications were accepted if they could be established at greater than 99.0% probability to achieve an FDR less than 1.0% and contained at least 2 identified peptides. Protein probabilities were assigned by the Protein Prophet algorithm[28].

### Whole Blood Transcriptomics and Weighted Gene Network Correlation Analysis

RNA-Seq was performed through the Accelerating Medicines Partnership-Parkinson’s Disease (AMP-PD) pipeline. RNA samples were rRNA and globin depleted via the Globin-Zero Gold kit (Illumina, USA) and first-strand synthesis and second strand synthesis were performed using the Ultra II First Strand Module and Ultra II Directional Second Strand Module (New England Biolabs, USA). Following second strand synthesis, the double-stranded cDNA was converted to a sequencing library by standard, ligation-based library preparation (see https://amp-pd.org/transcriptomics-data). Following ligation to standard paired-end adaptors (Illumina, USA), each library was amplified for 12 cycles using Kapa HiFi polymerase (Roche, USA), with forward and reverse primers including a 8 nucleotide unique index sequence. Insert sizes were evaluated via Caliper GX (Perkin-Elmer, USA) and libraries quantitated with Kapa SYBR FAST Universal kit (Roche, USA). Sequencing was performed on the Illumina NovaSeq 6000 platform to generate 100M paired reads per sample at 150 nucleotide read lengths. Samples were demultiplexed based on the unique i5 and i7 indices to individual sample FASTQ files. Only participants with full featureCounts were included here which was n=522. The featureCounts matrix was filtered to include all genes with non-zero transcript counts in greater than 25% of samples (n=36,168). A variance stabilizing transformation (VST) was applied using the DESeq2 V1.40.2 package[29]. DESeq2 was also utilized for differential analysis of count data in PD versus control subjects to identify differentially expressed genes (DEGs), where Benjamini-Hochberg corrected p values of <0.05 are considered significant.

The “WGCNA” R package was utilized to construct networks of coexpressed genes, where each node corresponds to a gene expression profile, and edges are determined by pairwise correlations between gene expression profiles across samples. A co-expression similarity matrix was constructed by calculating correlations among significant genes, which was converted to the adjacency matrix using a thresholding procedure. The optimal power coefficient β was identified by the R function pickSoftThreshold. A topological overlap matrix (TOM) was generated by the adjacency matrix, representing the strength of gene-gene interactions.

Alongside hierarchical gene clustering trees, the TOM identifies large modules connected to different colors. Module eigengenes (i.e. a weight average expression profile) were determined from the first principal component of each module, and significant modules were selected from module-trait relationships between eigengenes and pT73-Rab10 to total Rab10 ratios (adjusted p<0.05 considered significant). This analysis was repeated for LRRK2 serum levels and MDS-UPDRS III scores. Predicted protein-protein interaction networks in the modules were generated with StringDB (http://string-db.org). K-means clustering (n=10 clusters) was performed and clusters with the highest average local clustering coefficient were visualized. Reactome pathway ontological terms for the top cluster were also generated using StringDB.

### Statistical analysis software

GraphPad Prism 10, JMP Pro 16, and R-4.3.1 were utilized for all statistical tests. Types of statistical analyses used are indicated in Figure legends in addition to the numbers and types of independent observations that compose each experiment. All raw data associated with the figures are available online as Additional File 3.

## Results

### Development of sensitive and specific scalable assays for the measurements of the ratio of pT73-Rab10 to total Rab10 and LRRK2 in serum

Recent single molecule array (SiMOA) assays have been developed that successfully measure, to femtomolar levels, pathology biomarkers in neurodegenerative diseases from low volumes of biobanked serum or plasma[30]. Scalable assays for the measurement of the ratio of pT73-Rab10 to total Rab10 in biobanked biofluids have not previously been described. To fill this gap, we first screened efficiencies of different monoclonal antibody pairs for the detection of recombinant LRRK2 and Rab10 proteins in the SiMOA format. We, and others, have previously defined specificity of these antibodies in western blot formats using lysates from knockout or knockdown of LRRK2 and/or Rab proteins[7,31]. With the selected antibody pairs indicated in Fig 1A,B, alignments of the antibody-recognized epitopes onto AlphaFold structures showed solvent-oriented positioning with sufficient space between epitopes to allow simultaneous binding. Using recombinant full-length proteins for Rab10 and LRRK2 (Fig. 1C), standard curves were developed in custom-formulated sample buffers (e.g., “sample diluent buffer”). As both LRRK2 and pT73-Rab10 interact at the membrane in complex, and these proteins may be associated inside of extracellular vesicles, sample buffers were supplemented with denaturants (i.e., sodium deoxycholate) and reducing agents (i.e., dithiothreitol) not typically included in single-molecule assays in attempts to dissociate membranes and complexes while still allowing antibody binding to the desired epitopes. In the denaturing sample diluent buffer matrix, we could achieve sensitivity for the detection of recombinant Rab10 and LRRK2 proteins that reached low picogram per milliliter (Fig. 1D,E).

**Fig. 1.**
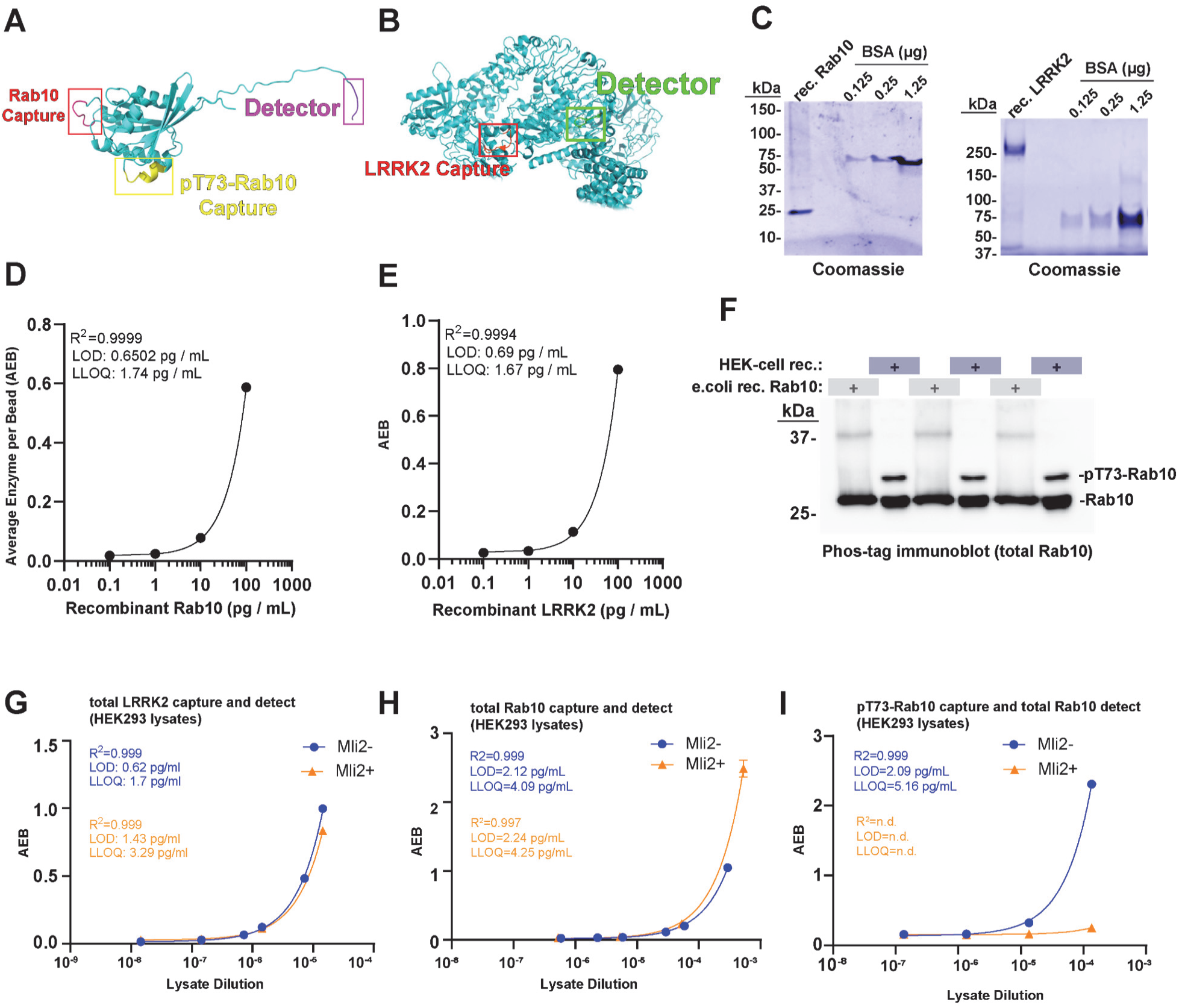
Development of sensitive single-molecule array assays and standards for fluid measures of total LRRK2, total Rab10, and pT73-Rab10. **(A)** AlphaFold structure of Rab10 and (**B)** LRRK2. Specific epitopes selected for the quantification of total LRRK2, total Rab10, and pT73-Rab10. The assay for Rab10 and pT73-Rab10 share the same detector antibody on the C-terminus but differ in capture antibody epitopes. **(C)** Representative SDS-PAGE gels with Coomassie stain (blue) for the quantification of recombinant His-Rab10 (derived from E. coli cells) and FLAG-G2019S-LRRK2 (derived from HEK-293T cells). Recombinant bovine-serum albumin standards are shown. **(D)** Representative regression curves for the measurement of recombinant Rab10 and (**E)** recombinant LRRK2 in the assay buffer. Goodness-of-fit (r-squared), limit-of-detection (LOD), and lower-limit of quantification (LLOQ) are indicated and calculated from duplicate and triplicate values for each assay point. **(F)** Representative phos-tag immunoblot analysis showing the proportion of Rab10 protein phosphorylated in protein lysates. HEK-293T lysates were generated through transient transfection of plasmids expressing FLAG-R1441G-LRRK2 with human FLAG-Rab10. The ratio of pT73-Rab10 to total Rab is calculated from three independent preparations as 15.2% ± 0.9% SEM. Protein preparations analyzed by phos-tag analysis are utilized as standards for single-molecule arrays (SiMOA). **(G)** Representative regression curves for the analysis of LRRK2, **(H)** Rab10, and (**I)** pT73-Rab10, as measured in diluted (sample buffer) HEK-293T protein lysates. Prior to lysis, cells were treated with MLi2 (200 nM, 30 min, red dots and lines) or vehicle (blue dots and lines). Assay quality parameters (orange font) are indicated based on triplicate values. Error bars for experimental replicates are typically less than 5% and too short to be visualized graphically in most of the plots.

As recombinant proteins may not mimic realistic matrix conditions (e.g., cell lysates, or viscous serum samples), LRRK2-transfected cell lines were processed and analyzed to determine the performance of the assay in the complexity of total protein lysates. First, a phos-tag approach was used to resolve the proportion of Rab10 phosphorylated according to western blot analysis of HEK-293T cells transfected with both FLAG-R1441G-LRRK2 with human FLAG-Rab10. Similar to proportions measured in previous reports[17,32], 15.2±0.9% of total Rab10 was found phosphorylated in the lysate (Fig. 1F). The SiMOA assays reported the concentration of LRRK2 protein and total Rab10 protein at 14.48±0.67 µg per mL for LRRK2 and 5.88±1.52 µg per mL (SEM) for total Rab10 in the lysates (Fig. 1G,H). Based on the measured proportion of pT73-Rab10 relative to non-phospho-Rab10 protein, the concentration of pT73-Rab10 in the lysate standard measured 0.89±0.23 µg per mL. Lower limits of detection and quantification in the lysates were slightly worse than those calculated for recombinant proteins (Fig. 1G-I). The analysis of pT73-Rab10 levels in HEK293T cells treated with the LRRK2 kinase inhibitor MLi2 prior to lysis shows that the pT73-Rab10 assay does not significantly cross react with non-phosphorylated Rab10 protein, even with high Rab10 levels in the lysates (Fig. 1I).

To determine whether these assays are viable in serum, known picogram quantities of recombinant LRRK2 and Rab10 were spiked into human serum samples and analyzed for recovered protein (after background subtraction). Both LRRK2 and Rab10 were recovered to ∼75% in serum, consistent with good assay performance (Supplemental Fig. 1A,B). A spike-in experiment using serum procured from *Lrrk2* knockout mice showed similar recovery of detection (Supplemental Fig. 1C). Higher signal (average enzyme per bead, or AEB) in the human serum samples that lacked any added recombinant proteins suggested the presence of detectable LRRK2 and Rab10 in serum.

To establish specificity of the assay signals in serum, LRRK2, Rab10 and pT73-Rab10 proteins were measured from low-volume single facial vein draws (∼40-60 µL) from a cohort of wild-type C57BL6/J mice, *Lrrk2^-/-^ (*knockout) mice, or *Lrrk2^R1441C/R1441C^* knockin mice (Fig 2A). Serum LRRK2 protein levels could not be determined in the *Lrrk2* knockout mice (Fig 2A), but were detectable in the other strains, suggesting good specificity for the LRRK2 assay. Serum Rab10 levels were similar between strains (Fig. 2B), demonstrating LRRK2 does not significantly regulate extracellular concentrations of total Rab10 levels in serum. pT73-Rab10 levels in LRRK2 knockout mice could not be determined, whereas the mean of the ratio of pT73-Rab10 to total Rab10 was low (0.79%±0.08%) in non-transgenic C57BL6/J mice. Notably, three of nineteen wild-type C57BL6/J mice had ratios of pT73-Rab10 to total Rab10 that were below the measured assay lower-limits of quantification (0.002, Figure 2C). However, the ratio of pT73-Rab10 to total Rab10 was measurable in all *Lrrk2^R1441C/R1441C^* knockin mouse serum samples, and, on average, was approximately twice that of wild-type C57BL6/J mice (Fig. 2C). These results suggest that wild-type mice have a very low basal state of phosphorylated Rab10 in serum, sometimes undetectable, where less than 1% of total Rab10 protein available is phosphorylated. Measured ratios in both the C57BL6/J and *Lrrk2^R1441C/R1441C^* mice were variable but appeared to be normally distributed (Fig 2C).

**Fig. 2.**
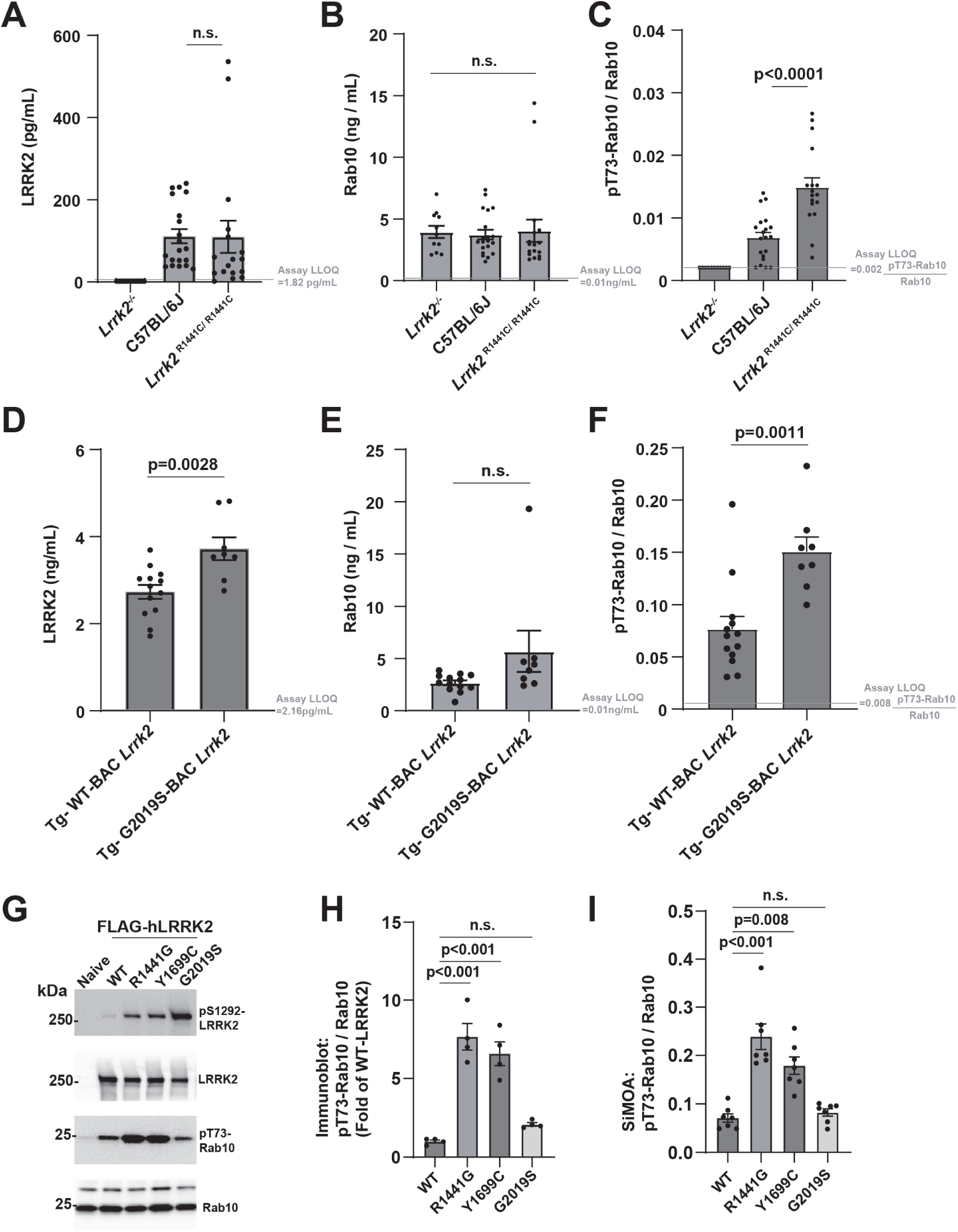
*LRRK2* mutations increase the extracellular ratio of pT73-Rab10 to total Rab10 as measured with single-molecule array assays (SiMOA) in serum. **(A)** Levels of total LRRK2, (**B)** total Rab10 and (**C)** the ratio of pT73-Rab10 to total Rab10 as measured by SiMOA in venous mouse serum from *Lrrk2*-knockout (8 males and 3 females), non-transgenic C57BL6/J (12 males and 8 females), and knockin *Lrrk2^R1441C/R1441C^* mice (9 males and 8 females). **(D-F)** A cohort of transgenic Tg-WT-BAC *Lrrk2* (4 males and 9 females) and Tg-G2019S-BAC *Lrrk2* mice (5 males and 3 females) were also analyzed for these markers. In these strains, no differences were noted between males and females. Ages in both cohorts of the mice ranged from 3-6 months. **(G)** Representative immunoblots of lysates from HEK-293T cells transfected with FLAG-LRRK2 plasmids 24-hours before harvesting lysates. **(H)** Quantification of endogenous pT73-Rab10 to total Rab10 ratio. **(I)** SiMOA assay analysis of the lysates for the ratio of pT73-Rab10 to total Rab10. Dots show mean values calculated from independent experiments. Columns show group means, error bars show SEM, and p values are determined with two-tailed t-tests, or from one-way ANOVA followed by Tukey’s post-hoc test in the graphs with more than two groups.

The effects of the pathogenic G2019S-*LRRK2* mutation on pT73-Rab10 levels has been controversial, with some studies showing G2019S-LRRK2 does not increase phosphorylated Rab10 levels over WT-LRRK2[16,17,21]. Our previous work measuring LRRK2 and pT73-Rab10 proteins using western blots demonstrated that peripheral macrophages isolated from transgenic mouse-BAC Tg-WT-*Lrrk2* and Tg-G2019S-BAC *Lrrk2* mice have similar levels of total LRRK2 protein compared to peripheral macrophages isolated from human blood, with non-transgenic mice and *Lrrk2* knockin mice showing much lower levels of LRRK2 protein, irrespective of interferon-stimulation of *LRRK2*/*Lrrk2* expression[12]. Total Rab10 levels seem independent from LRRK2 as LRRK2 knockouts have normal Rab10 levels. Analysis of serum from Tg-G2019S-BAC *Lrrk2* mice show, on average, increased extracellular LRRK2 protein compared to Tg-WT-BAC *Lrrk2* mice (Fig. 2D), but similar levels of total Rab10 (Fig. 2E). Serum from Tg-G2019S-BAC *Lrrk2* mice shows much higher ratios of pT73-Rab10 to total Rab compared to wild-type or *Lrrk2* knockin mice, averaging 15.1±1.4% of total Rab10 phosphorylated (Fig. 2F). In comparison to Tg-G2019S-BAC *Lrrk2* serum, Tg-WT-BAC *Lrrk2* serum shows about half the amount of phosphorylated Rab10 on average (7.7±1.2%). These results show extracellular pT73-Rab10 is sensitive to LRRK2 expression and the G2019S-*LRRK2* mutation.

To further explore the effects of different pathogenic *LRRK2* mutations on the ratio of pT73-Rab10 to total Rab10, as measured by the SiMOA assay in cell lysates, HEK-293T cells were transfected with FLAG-WT-LRRK2, R144G, Y1699C, and G2019S-LRRK2, and verified for pT73-Rab10 levels with standard western blot analysis (Fig. 2G, H). The activating effects of G2019S-LRRK2 on autophosphorylated pS1292-LRRK2 were also confirmed by western blot. The SiMOA assay measured 23.9±2.6% and 17.9±1.8% phosphorylated Rab10 for R1441G and Y1699C-LRRK2, respectively, in comparison to 7.1±0.9% for WT-LRRK2 (Fig. 2I). However, G2019S-LRRK2 expression in cell lysates did not increase pT73-Rab10 to total Rab10 ratios over those measured with WT-LRRK2 expression.

### The extracellular ratio of pT73-Rab10 to total Rab10 primarily derives from bone-marrow immune cells and increases with inflammation

According to recent transcriptomic studies, the highest *LRRK2* mRNA levels in cells and tissues across the body may reside in myeloid cell populations, particularly neutrophils, monocytes and macrophages, and microglia[12,33,34]. Circulating myeloid cells can be efficiently ablated with a single whole body irradiation procedure (Fig. 3A). In CD-1 outbred mice, six days after irradiation, there were few leukocytes left in blood. In these irradiated mice, LRRK2 protein in serum drops to near undetectable levels (Fig. 3B), and both total Rab10 as well as the ratio of pT73-Rab10 to total Rab10 drops considerably compared to baseline (Fig. 3C,D). Transplantation of bone-marrow cells back to the irradiated mice re-establishes leukocyte population counts in the blood (Fig. 3E). In the transplanted mice, normal serum levels of LRRK2 and the ratio of pT73-Rab10 to total Rab10 were measured (Fig. 3F-H). These results suggest that the ratio of pT73-Rab10 to total Rab10 in serum primarily originates from bone-marrow derived cells.

**Fig. 3.**
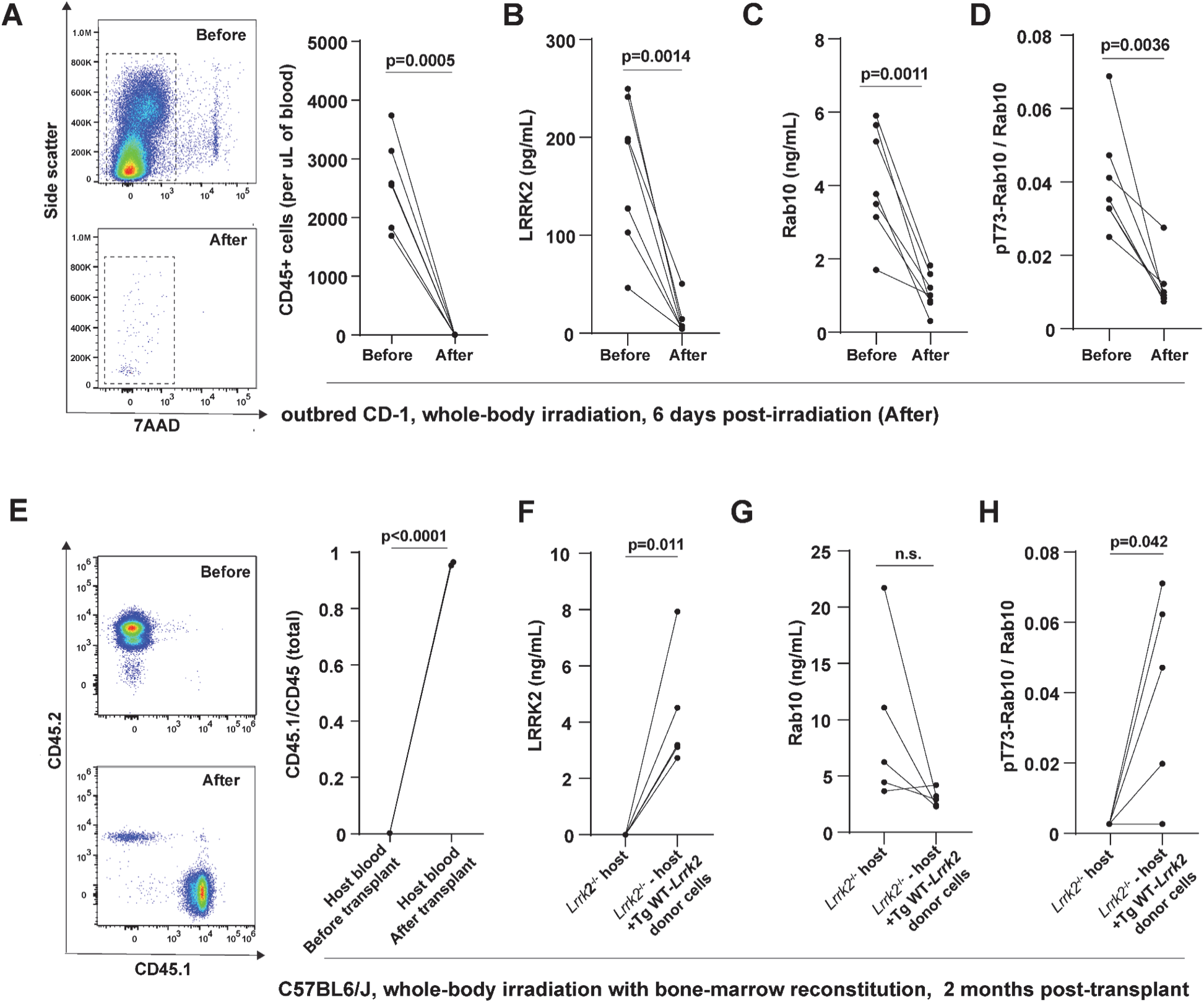
Serum LRRK2 and pT73-Rab10 proteins originate primarily from bone marrow immune cells. **(A)** Representative cytographs and column graphs showing the analysis of live (7AAD-7-aminoactinomycin D negative) leukocytes in blood from outbred male CD-1 mice before and six-days after 11 grays of radiation. **(B)** SiMOA analysis of serum from these mice for **(B)** total LRRK2, (**C)** total Rab10 and (**D)** the ratio of pT73-Rab10 to total Rab10. **(E)** Cytographs show the analysis of host CD45.2 and donor CD45.1 mice before and after radiation, and the column graph shows the mean of the ratio of CD45.1 cells to total CD45 cells (n=3 male mice). Bone marrow transplantation of donor Tg-WT-*Lrrk2* cells into host *Lrrk2^-/-^* mice restores levels of **(F)** total LRRK2, (**G)** total Rab10 and (**H)** the ratio of pT73-Rab10 to total Rab10. Each dot shows mean values from the analysis of a single animal (all ∼2 months of age), and p values are calculated from paired (before and after) sample t-tests.

Our and other previous work suggests LRRK2 expression may increase in myeloid cells after exposures to lipopolysaccharide or interferon-induction as part of a pro-inflammatory response[35,36]. The role of phosphorylated Rabs such as pT73-Rab10 in these responses is not yet clear, though our recent studies in macrophages implicate pT73-Rab10 in stalling signaling endosomes from turnover in canonical AKT pathways[7]. To test whether acute and systemic inflammation in mice might affect extracellular pT73-Rab10 to total Rab10 ratios, we introduced a polymicrobial sepsis challenge to outbred CD-1 mice that caused large increases in serum concentrations of IFNγ and TNF (Fig. 4A-C). Surprisingly, 72-hours post sepsis, LRRK2 levels only modestly increase in female mice, and not at all in male mice (Fig. 4D). As IFNγ is thought to be a principal driver of LRRK2 expression in myeloid cells[37], higher IFNγ levels that develop in female mice may explain higher extracellular LRRK2 levels. Further, sepsis drops the levels of total Rab10 levels in serum (Fig 4E). However, the measured ratio of pT73-Rab10 to total Rab10 increases in both male and female mice (Fig. 4F). These results show that the ratio of pT73-Rab10 to total Rab10 in serum increases with sepsis, albeit with variability between animals. One each of the wild-type male and female mice developed more than 20% phosphorylated Rab10 ratios in serum, exceeding the measured pT73-Rab10 to total Rab10 ratios measured in all of the Tg-WT-BAC *Lrrk2* under basal conditions (Fig 2F). The non-transgenic mice achieve this level of pT73-Rab10 phosphorylation despite expressing 10-fold lower levels of LRRK2 protein as measured in the serum (0.21±0.02 vs. 2.7±0.16 ng/mL in the Tg-WT-BAC *Lrrk2* mice, males and females combined). These results show that LRRK2 protein is required, but not sufficient, for elevated extracellular pT73-Rab10 to total Rab10 ratios, and that wild-type mice can develop high levels (albeit variable) of phosphorylated Rab10 protein in serum.

**Fig. 4.**
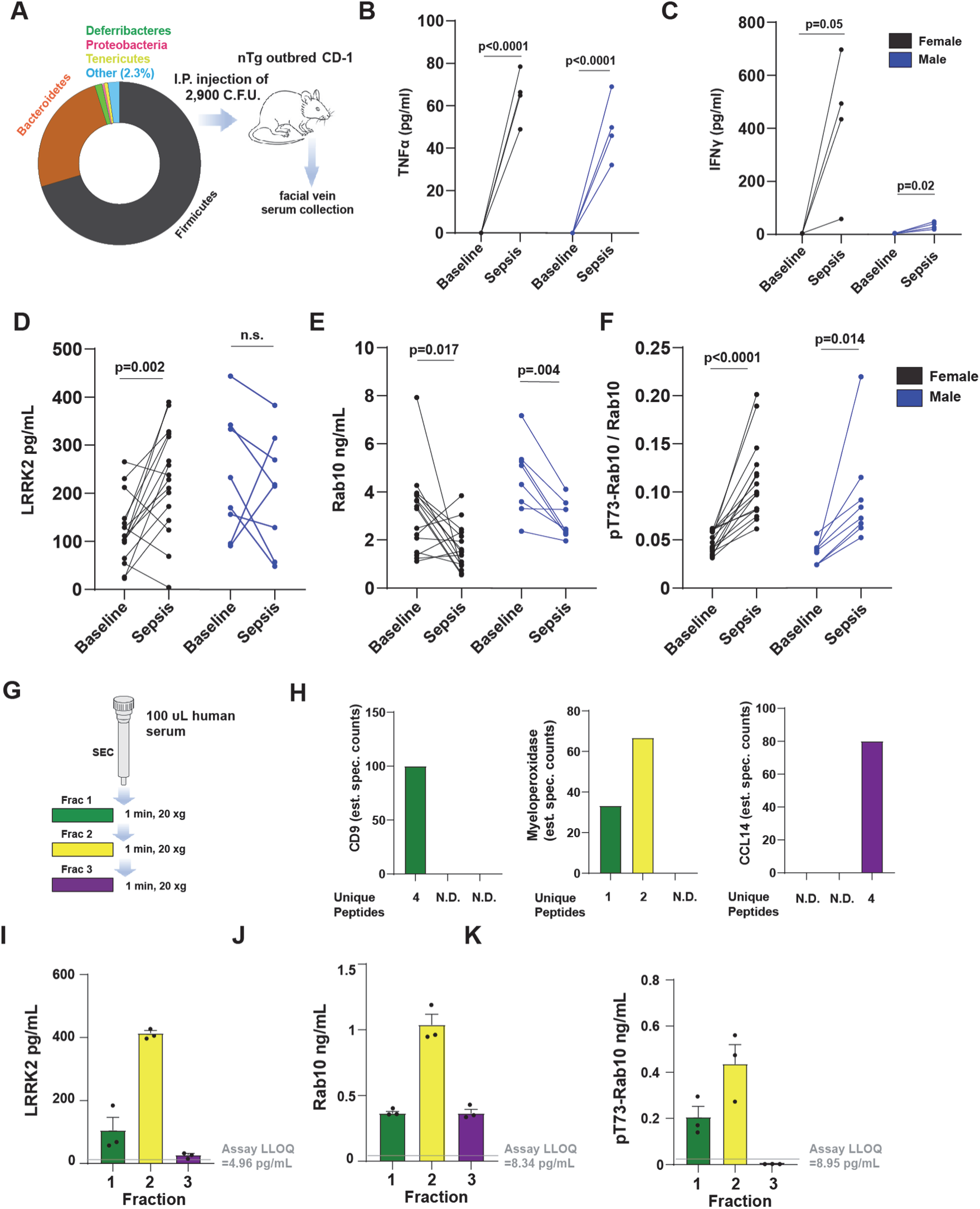
Sepsis increases the extracellular ratio of pT73-Rab10 to total Rab10 in serum that primarily localizes to exosome-enriched serum fractions. **(A)** Microbe profile of cecal slurry generated from healthy CD-1 outbred male mice. Percentages of phylum-level taxonomy, identified by 16S rRNA amplicon sequencing, are depicted with the “Other” category including *Actinobacteria, Cyanobacteria,* and *Verrocomicrobia.* Slurries were injected intraperitoneally into a cohort of outbred male (n=8) and female (n=16) CD-1 mice, with serum procured from facial vein draws. At the 8 hr time point after injection, 4 female and 4 male mice were selected at random to measure serum **(B)** interferon-γ (IFNγ) and **(C)** TNF with ELISA analysis. **(D)** Before and after graphs show changes in serum LRRK2 levels, **(E)** total Rab10 and **(F)** the ratio of pT73-Rab10 to total Rab10 72 hrs post-injection. P values shown are from paired samples (before and after) t-tests. **(G)** Procedure for “Exo-Spin Serum mini columns’’ size-exclusion fractionation of 100 microliters of human serum spread into fractions 1-3. **(H)** Proteomic analysis of eluted fractions via mass spectrometry assessments of the relative abundance of unique peptides associated with characteristic serum exosome proteins CD9 (plasma-membrane protein enriched in exosomes and extracellular vesicles), myeloperoxidase (a soluble myeloid-cell produced enzyme enriched in exosomes and extracellular vesicles), and a small soluble cytokine HCC-1 (CCL14). **(I)** Column graphs show levels of LRRK2, **(J)** Rab10, **(K)** and pT73-Rab10 as measured in three different human biobanked serum samples (one male and one female with PD, and one healthy control). Green bars are measured from fraction one, yellow bars are fraction two, and purple are fraction three. Columns show group mean and error bars show SEM.

Myeloid cells are well known to secrete small EVs as well as shed matrix proteins into the extracellular milieu to control immunological responses. Myeloid cells secrete proinflammatory EVs that include exosomes in response to immunological challenges, and both LRRK2 and Rab10 have been detected in extracellular protein fractions enriched in EVs[38]. Separation of human serum (n=3 human serum samples) through a size-exclusion column optimized for serum separation of EVs from other extracellular compartments yielded three fractions that we characterized by whole-proteome analysis (Fig. 4G). The first elution, containing larger complexes, serum EVs and exosomes, shows peptides from canonical exosome proteins including CD9 (Fig. 4H). The pro-inflammatory myeloperoxidase protein, known to be upregulated with inflammation in myeloid cells and subsequently shed at high levels in both exosomes and non-EV serum fractions[39,40], appears split in concentration between the first two elutions. Cytokines and chemokines of low molecular weight were present in the third elution (Fig 4H). Similar to the distribution of myeloperoxidase, LRRK2 and pT73-Rab10 were both confined to fractions 1 and 2 from the column (Fig 4I,J), whereas total Rab10 was also detected in fraction 3 (Fig. 4K). These results suggest that both LRRK2 and pT73-Rab10 may share a similar extracellular compartment in serum, enriched in EVs, but may not exclusively localize to exosomes.

### The serum ratios of pT73-Rab10 to total Rab10 are elevated in *VPS35* and *SNCA* mutant mice

Recently, a pathogenic *VPS35* mutation, D620N, has been described to increase cellular levels of pT73-Rab10 in tissue lysates in a LRRK2-kinase dependent manner[41]. In serum samples procured from D620N-*VPS35* knockin mice, total LRRK2 and Rab10 protein concentrations are similar between littermate WT, heterozygous, and homozygous mice (Fig. 5A,B). In contrast, the ratio of pT73-Rab10 to total Rab10 is elevated in *VPS35*^D620N/D620N^ mice to 7.1±0.7% (Fig 5C), similar to the average ratios in Tg-WT-BAC *Lrrk2* serum with over-expressed *Lrrk2* (7.7±1.2%, Fig. 2F) as well as many of the septic wild-type mice (Fig. 4F). Analysis of whole blood from *VPS35*^D620N/D620N^ mice compared to wild-types shows the *VPS* mutant mice may have increased neutrophils and CD4-positive T cells, but similar numbers of monocytes (Fig. 5D-F). Western blot analysis of lysates derived from cultured bone-marrow derived macrophages (BMDMs) from the *VPS* mutant mice highlight the expected activating effects of the *VPS35* D620N mutation on increasing the normalized (to wild-type mice) ratio of pT73-Rab10 to total Rab10 (Fig 5G). Interestingly, pT73-Rab10 was not reliably detected by either SiMOA or western blot in litter-mate wild-type mice processed at the same time. These results suggest that the *VPS35* mutation increases serum levels of the ratio of pT73-Rab10 to total Rab10 without affecting serum LRRK2 or total Rab10 levels.

**Fig 5.**
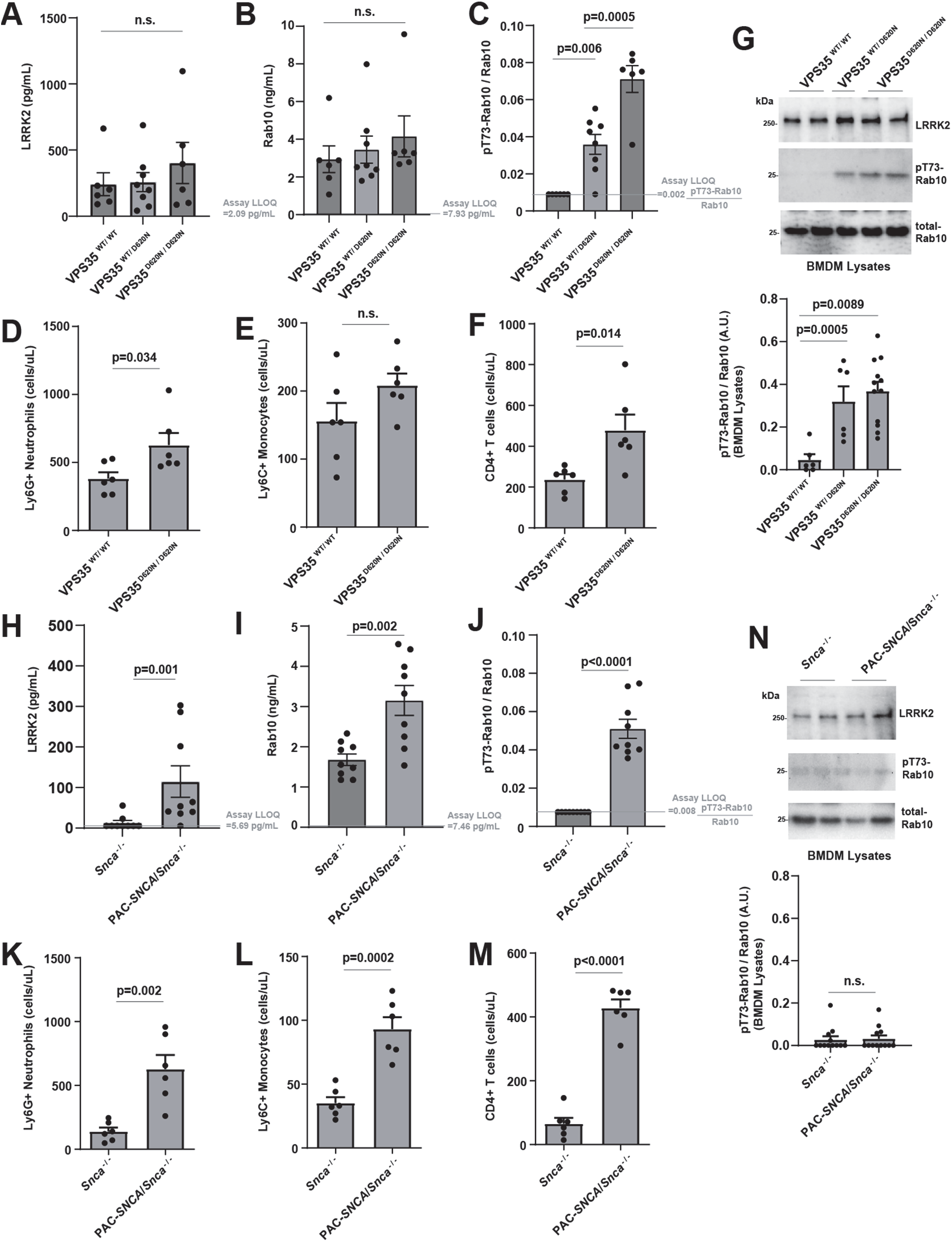
The ratio of pT73-Rab10 to total Rab10 in serum is elevated in mouse models of late-onset PD. **(A)** Levels of total LRRK2, (**B)** total Rab10 and (**C)** the ratio of pT73-Rab10 to total Rab10 as measured by SiMOA in venous serum from *VPS35^WT/WT^* (2 males and 4 females) mice, *VPS35^WT/D620N^* (4 males and 4 females) mice, and *VPS35^D620N/D620N^* (6 males) mice. **(D)** Graphs show numbers of neutrophils (CD11b+ Ly6G+), **(E)** classical monocytes (CD11b+ Ly6G-Ly6C+), and **(F)** CD4 T-cells measured by flow cytometry in *VPS35^WT/WT^* and *VPS35^D620N/D620N^* serum samples. **(G)** Representative immunoblots and quantification of cultured bone marrow derived macrophages demonstrating elevated pT73-Rab10 to total Rab10 levels with mutant VPS35 expression. **(H)** Levels of total LRRK2, (**I)** total Rab10 and (**J)** the ratio of pT73-Rab10 to total Rab10 from *Snca^-/-^* (3 males and 6 females) mice, and PAC*-SNCA/Snca^-/-^* (5 males and 4 females) mice. **(K)** Graphs show numbers of neutrophils (CD11b+ Ly6G+), **(L)** classical monocytes (CD11b+ Ly6G-Ly6C+), and **(M)** CD4 T-cells measured by flow cytometry in *Snca^-/-^* and PAC*-SNCA/Snca^-/-^* mice blood. **(N)** Representative immunoblots and quantification of cultured bone marrow-derived macrophages demonstrating low but equivalent pT73-Rab10 to total Rab10 between *Snca^-/-^* and PAC*-SNCA/Snca^-/-^* mice. Mouse ages ranged from 3-6 months. Columns depict group mean, dots represent the mean values from an animal or independent experiment, and error bars show SEM. P values are from two-tailed t-tests or one-way ANOVA analysis with Tukey’s multiple comparison when three groups are tested.

We recently reported pathological α-synuclein aggregates may stimulate levels of pT73-Rab10 in a LRRK2-activity dependent manner in myeloid cells[12]. A transgenic mouse line expressing wild-type human α-synuclein (PAC-*SNCA*/*Snca^-/-^* mice) harbors significant concentrations of α-synuclein aggregation and seeding activity according to past reports, accompanied by pronounced gastrointestinal disturbance phenotypes[42,43]. In serum from these mice, both LRRK2 protein and total Rab10 is significantly increased compared to *Snca^-/-^* control mice (Fig 5H,I). The elevated ratio of pT73-Rab10 to total Rab at 5.1%±0.5 in serum from these mice, higher than levels found in R1441C-*LRRK2* homozygous knockin mice, suggests the possibility of inflammation (Fig 5J). Flow cytometry analysis of whole blood from the same mice confirmed upregulated neutrophil populations, monocytes, as well as T-cells (Fig. 5K-M). However, different from the *VPS35* mutant mice, western blot analysis of isolated and cultured BMDM cells from the the PAC-*SNCA*/*Snca^-/-^* mice did not show intrinsically upregulated pT73-Rab10 levels compared to controls (Fig 5N). Taken together, these data suggest that the extracellular ratios of pT73-Rab10 to total Rab10 in serum can be sensitive to genetic variants that stimulate LRRK2 activity (e.g., R1441C-*Lrrk2* or *VPS35* mutation), acute inflammation (e.g., sepsis), as well as chronic inflammation found in a progressive iPD α-synuclein transgenic model. Under basal conditions, serum pT73-Rab10 to total Rab10 ratios are very low in mice, undetectable in some cases. In all of these mouse cohorts evaluated at baseline, there were no differences observed between male and female mice from the same strain in serum LRRK2, Rab10, and pT73-Rab10 levels.

### The ratio of pT73-Rab10 to total Rab10 in serum is pharmacodynamic

Cellular levels of pT73-Rab10 are known to be highly pharmacodynamic and drop within minutes of LRRK2 kinase inhibition[44], but the extracellular ratio of pT73-Rab10 to total Rab10 has not been previously measured in response to drugs. The small molecule LRRK2 inhibitor PFE-360 is orally available and quickly metabolized in mice and rats, with a serum half life in oral dosing of t½= ∼1.3 hours in rats and a potency of ∼IC_50_ 6 nM *in vitro* for LRRK2 kinase inhibition[45]. To determine the pharmacodynamic potential of the extracellular ratio of pT73-Rab10 to total Rab10 in serum, we produced longitudinal sets of serum samples from rodent strains with elevated (i.e., more than wild-type mice) pT73-Rab10 levels, Tg-G2019S-BAC *Lrrk2* and *VPS35* mutant mice, treated with a single dose (10 mg per kg) of PFE-360. Measured serum levels of the ratio of pT73-Rab10 to total Rab10 are depressed within 30 minutes of the oral gavage of drug in both strains of mice (Fig. 6A,B). Still lower levels were recorded at one-hour post dose, 78.3±9.1% inhibition in *VPS35* knockin mice and 83.3±7.4% inhibition in Tg-G2019S-BAC *Lrrk2* mice. In the facial vein collections of serum, when insufficient volume or a hemolyzed blood sample was collected, the sample was omitted from analysis and collection attempted again at the next time point (Fig 6A-C). At one hour post-drug treatment in wild-type rats, serum levels of the ratio of pT73-Rab10 to total Rab10 were only 37.4±8.3% lower compared to baseline levels, with continued suppression evident 24 hours post-gavage (Fig. 6C).

**Fig. 6.**
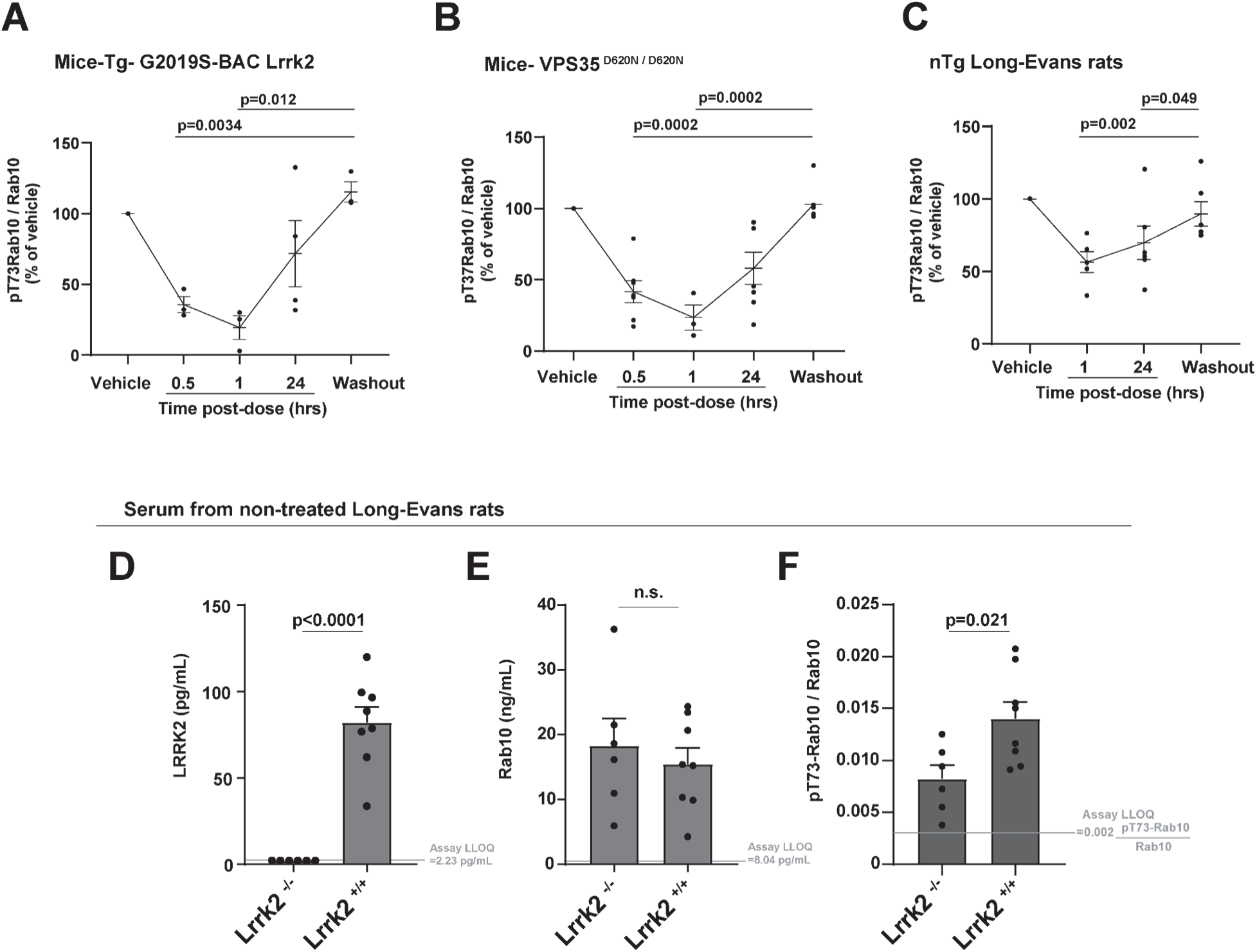
The ratio of pT73-Rab10 to total Rab10 in serum is pharmacodynamic. In mice and rats, the LRRK2 small molecule inhibitor PFE-360 was given as a single oral dose (10 mg per kg), serum (mouse facial vein or rat lateral saphenous vein) collected one-hour after a vehicle-only administration (Vehicle), and serum next collected at the indicated time post-drug administration, with a final collection at 48 hours after drug (Washout). The ratio of pT73-Rab10 to total Rab10 diminishes relative to baseline in **(A)** Tg-G2019S-BAC-*Lrrk2* (4 male mice) and **(B)** *VPS35^D620N/D620N^* (7 male mice), as well as **(C)** nTg Long-Evans *Lrrk2^+/+^* rats (7 male rats). Rodent ages were 3-5 months, bars show group means, error bars show SEM, and dots show the mean values from individual rodents. In repetitive venous draws, some samples were not successfully collected due to insufficient volume collected or hemolysis, thereby excluding the sample for analysis (see Additional File 3 for all raw data collected). Bars in panels A-C show group mean, error bars show SEM, and p values were calculated from one-way ANOVA analysis with Tukey’s post-hoc comparisons. **(D)** Graphs show levels of LRRK2, **(E)** Rab10, and **(F)** the ratio of pT73-Rab10 to total Rab10 in Long-Evans *Lrrk2^-/-^* rats (3 males and 3 females) compared to non-transgenic Long-Evans *Lrrk2^+/+^* rats (4 males and 4 females). Columns in panels D-E show group mean, error bars show SEM, and p values are from two-tailed t-test.

To pinpoint why only a minority of pT73-Rab10 was inhibited in the wild-type rats, we generated another set of serum samples from a new cohort of wild-type (Long Evans outbred) rats together with serum from *Lrrk2* knockout rats (also Longs Evans background), with the knockout rat serum representing the lowest ratio of pT73-Rab10 to total Rab10 attributable to LRRK2. As anticipated, there were no measurable quantities of LRRK2 protein in serum from *Lrrk2* knockout rats (Fig. 6D). Serum levels of total Rab10 were similar between the wild-type and knockout strains (Fig. 6E). However, the ratio of pT73-Rab10 to total Rab10 was only partially lowered in the *Lrrk2* knockout rats compared to wild-types. The reduced level in the knockout rats was similar to the reduction measured one hour after PFE-360 gavage (41.1±8.7% reduction in the knockout compared to 37.4±8.2% reduction with PFE-360). These results suggest that serum pT73-Rab10 levels in rats, which are low to begin with (all rat samples measured <2.5% pT73-Rab10 to total Rab10), are only partially dependent on LRRK2 under basal conditions.

### Ratios of pT73-Rab10 to total Rab10 are elevated in PD cases with higher MDS-UPDRS scores

To better understand the ratio of pT73-Rab10 to total Rab10 in the periphery in iPD, we selected a cohort of serum samples donated from participants enrolled in the Parkinson’s Disease Biomarker Program (PDBP). 522 serum samples were available from 277 PD cases and 246 controls originating from a single collection site (Table I). Serum samples matched whole-blood transcriptomic and genetic data generated from participant tubes of blood collected at the same time as the serum samples, and all clinical and demographic data were collected during the same visit as the blood tube collections. Blood draws occurred between the hours of 8 and 10 in the morning, with all participants taking their normal routine of medications. L-dopa equivalent daily dosages (LEDD) at the time of sample donation were calculated. Clinical and demographic characteristics in the cohort highlight lower UPSIT scores in the PD group, reflecting hyposmia in some cases, and nominally worse cognition reflected in lower MoCA scores (Table I). The PD and control group had a similar usage of anti-inflammatories (NSAIDs) and statins. LEDDs are moderately correlated to motor defects reflected in MDS-UPDRS part III scores (r=0.436, p<0.0001).

SiMOA measurements of total LRRK2, total Rab10, and pT73-Rab10 in human sera were made without respect to group assignment in a randomized fashion from coded samples. Analysis of assay performance across the cohort shows a dynamic linear range of ∼2-3 orders of magnitude in sensitivity in each run, consistent lower limits of quantification (LLOQ) from run-to-run, and low variability (coefficient of variations) across technical replicates (Supplemental Fig. 2). Similar to studies in rodents, ∼60 microliters of serum were utilized for measurements of all three analytes with technical replicates. In addition to LRRK2, total Rab10, and pT73-Rab10, serum creatinine values were measured in the same samples by ELISA. Creatinine levels for all samples fell within normal ranges and did not correlate with total LRRK2 levels (r=-0.017, p=0.70), total Rab10 (r=0.069, p=0.12), pT73-Rab10 levels (r=0.024, p=0.58), or the ratio of pT73-Rab10 to total Rab10 (r=0.065, p=0.14). All samples were estimated to have hemoglobin concentrations less than 100 mg per dL using standard scales. Twenty serum samples, selected at random, were analyzed for exact hemoglobin values via ELISA analysis, and no correlations were found between hemoglobin levels and any of the other proteins measured. Further, no correlations were found between LRRK2, total Rab10, or pT73-Rab10 and freezer storage time.

In the analysis of total LRRK2 protein, mean values of LRRK2 at 6,537 pg per mL (95% C.I. [1,501 to 11,573]) across all human samples were, on average, much higher than those observed in serum from mice (111±17 pg per mL [SEM]) and rats (82±6 pg per mL [SEM]). However, the Tg-*Lrrk2* BAC mice (mean of both WT and G2019S strains) had LRRK2 expression (3,460±265 pg per mL [SEM]) closer to that found in humans. These results may be consistent with observations from western blots we previously performed from macrophage cell lysates, where LRRK2 protein levels were similar between BAC strains and human primary macrophages, and much lower in macrophages from non-transgenic mice, irrespective of interferon stimulation[12].

Male PD subjects had slightly higher LRRK2 expression than female controls, with an average difference of 0.96 ng per mL (95% C.I.[0.26 to 1.65], adjusted p=0.003, Supplemental Fig. 3). However, a least squares multivariate regression approach did not reveal any significant associations between LRRK2 levels in serum and clinical rating scales including MDS-UPDRS scores, MoCA, UPSIT, and Epworth Sleep scales, with all clinical scales recorded within hours of the serum sample donation in the on-medication state. Across the cohort, total Rab10 levels are also much higher in the average human serum sample (mean 78 ng per mL, 95% C.I. [56 to 100]) compared to the average serum sample from non-transgenic mice (4±0.5 ng per mL [SEM]) and Long-Evans rats (15±3 ng per mL [SEM]). Despite successful measurements of total LRRK2 and total Rab10 levels in all samples in the 522 sample cohort, a sizable proportion of samples (23.5%) had levels of pT73-Rab10 below the empirically determined average lower limit of quantification (LLOQ, 10.1 pg of pT73-Rab10 per mL, Supplemental Fig. 3). To prevent the assignment of a null value for the ratio of pT73-Rab10 to total Rab10 for these samples and exclusion from further statistical analysis, the lower limit value of 10.1 pg per mL for pT73-Rab10 was divided by the measured total Rab10 in the sample to obtain an overestimated numerical ratio for pT73-Rab10 to total Rab10. Because total Rab10 tends to be high in all human serum samples, the ratio value of pT73-Rab10 to total Rab10 in these samples remained very low, with a mean ratio of 0.0036 for pT73-Rab10 to total Rab10, 95% C.I. [0.0030 to 0.0042]. This low level of phosphorylation can be considered biologically insignificant since comparable ratios were measured in *Lrrk2* knockout mice.

In rodents, the highest recorded ratio of pT73-Rab10 to total Rab10 was 38.2%, found in a serum sample from a male Tg-G2019S-BAC *Lrrk2* mouse. While many human serum samples had very low pT73-Rab10 levels, comparable to levels observed in wild-type mice and rats under basal conditions, a sizable proportion of human serum samples (9.8%, or 51 of 522 samples) had ratio values that exceeded 38.2% (i.e., the highest recorded value observed in transgenic mice). Two of the 522 samples (0.4% of total samples) had a ratio of pT73-Rab10 to total Rab10 above 1.0, specifically, 1.25 and 1.27, possibly reflecting technical error associated with the regression of the assay at the very high end of the measures. Both these samples were adjusted to ratio values of 1.0 that represent the upper limit of the assay. The ratio of pT73-Rab10 to total Rab10 was variable across the cohort, and efforts to improve normalcy of the distribution with additional transformations were not successful (Supplemental Fig 3).

Distributions of the ratio of pT73-Rab10 to total Rab10 were overall similar between participants with and without PD (Supplemental Fig. 3). Within the cohort of samples, two control subjects donated serum weekly across a two month interval. LRRK2 serum levels and the ratio of pT73-Rab10 to total Rab10 were relatively stable over this time period, with one subject consistently showing higher LRRK2 and pT73-Rab10 to total Rab10 ratios than the other subject (Supplemental Fig. 4).

In samples across the cohort, serum LRRK2 protein levels strongly correlate with serum total Rab10, pT73-Rab10, and the ratio of pT73-Rab10 to total Rab10 (Fig. 7A-C). A stepwise regression analysis including all demographic and clinical variables listed in Table I, as well as LRRK2 genotypes G2019S, rs76904798, N2081D, as well as N551K/R1398H, found that MDS-UPDRS scores, as well the G2019S-*LRRK2* mutation and total LRRK2 levels significantly predicted the ratio of pT73-Rab10 to total Rab10. In a least squares model (RMSE=0.20, r=0.33, p<0.0001) that includes MDS-UPDRS part II scores, G2019S-*LRRK2* mutation status, and LRRK2 levels in serum, a doubling of LRRK2 serum levels is associated with an increase of 1.4% of pT73-Rab10 to total Rab10 (95% C.I. [0.40% to 0.24%], p=0.0063). An increase of a point on the MDS-UPDRS part III is associated with an increase of 0.28% pT73-Rab10 to total Rab10 (95% C.I. [0.089% to 0.47%], p=0.0043). Finally, the presence of the G2019S mutation is associated with a larger increase of 19.0% (95% C.I. [8.99% to 29.0%], p=0.0002). Other variables in the model including age, sex, disease duration, L-dopa equivalent daily dose (LEDD), and current usage of non-steroidal anti-inflammatories (NSAIDs), did not significantly predict the ratio of pT73-Rab10 to total Rab10.

**Fig. 7.**
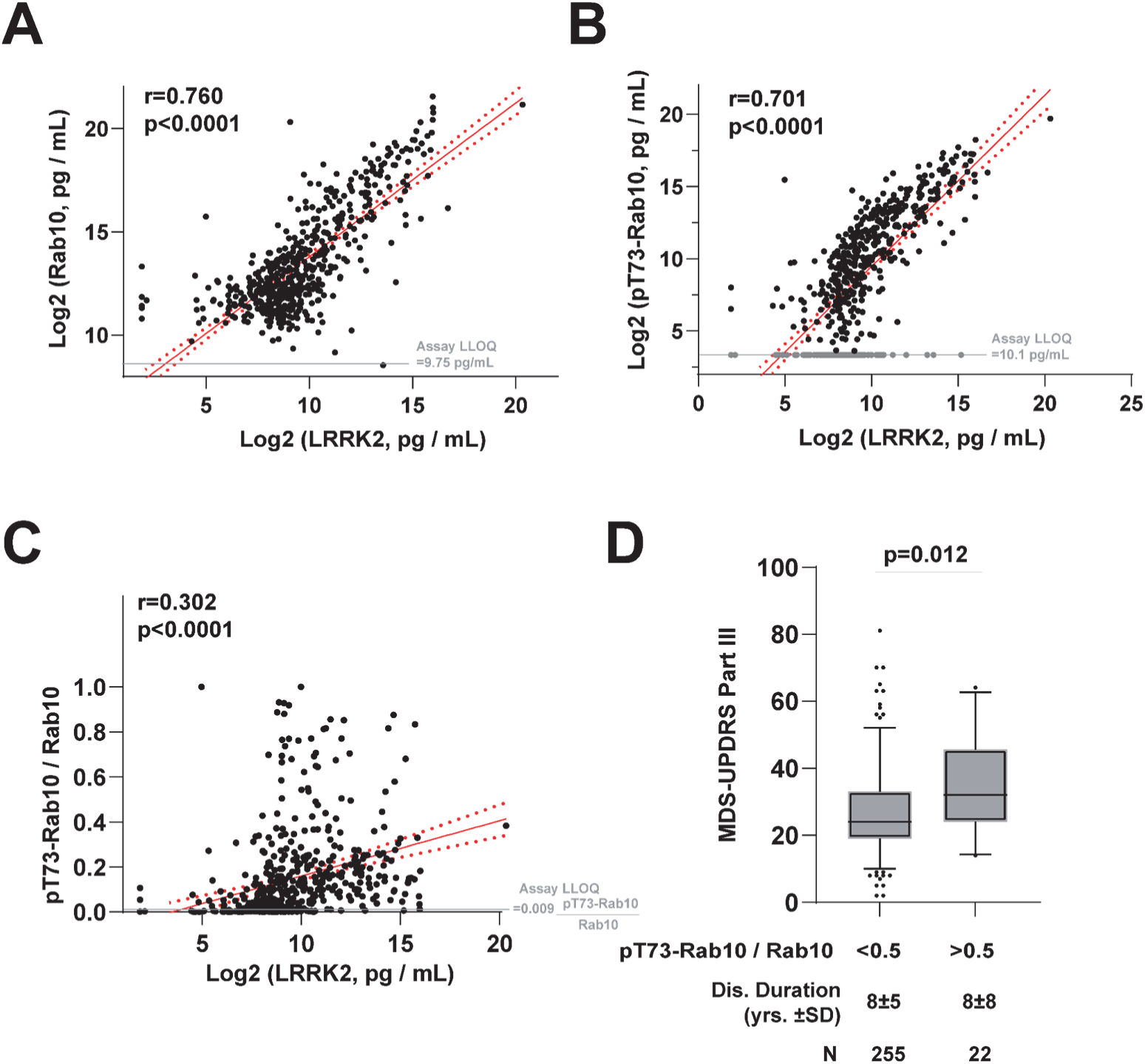
LRRK2 protein levels predict pT73-Rab10 levels that are elevated in PD cases with worse MDS-UPDRS scores. **(A)** Scatter plots show relationships between measured serum levels of total Rab10 and LRRK2, **(B)** pT73-Rab10 and LRRK2, and **(C)** the ratio of pT73-Rab10 to Rab10 and LRRK2. N=522 cases and controls (see Table I). R values show correlation coefficients and p values correspond to mean linear regression analysis of lines (red) with 95% confidence intervals shown (red dashed curves). **(D)** Box and whisker plot comparing the mean MDS-UPDRS part III scores in PD cases split to lower (<0.5) and higher (>0.5) ratios of pT73-Rab10/Rab10. P value indicated is from a Mann-Whitney U test for comparisons of groups with unequal sizes.

While only four subjects in the cohort had a G2019S-*LRRK2* heterozygous mutation, all four had a very high measured serum ratio of pT73-Rab10 to total Rab10 (mean value 52.1±16.9% [SEM]). As the G2019S-*LRRK2* mutation is considered pathogenic in PD cases, a rough initial estimation of a pathogenic level of Rab phosphorylation may be represented through the average ratios found in the mutation carriers, or ∼50% pT73-Rab10 to total Rab10. A strata of iPD cases without *LRRK2* mutations demonstrated this level of pT73-Rab10 or higher. Splitting the PD cases in the cohort between participants with high (e.g., possibly pathogenic) pT73-Rab10 to total Rab10 ratios that exceed 50% from those with levels less than 50% highlights worse MDS-UPDRS part III scores in PD cases with the higher Rab10 phosphorylation (Fig. 7D). The higher pT73-Rab10 group is 7.3 points worse on the MDS-UPDRS part III score than those with lower Rab phosphorylation (p=0.012, Mann Whitney U test). The difference of 7.3 points on the MDS-UPDRS part III is associated with approximately three years of disease progression in iPD[46], but there is a similar disease duration between the high and low pT73-Rab10 groups (∼8 years on average, Fig. 7D), and there is no correlation between disease duration and the ratio of pT73-Rab10 to total Rab10 (r=-0.006, p=0.92). Splitting the cohort according to PD diagnosis, NSAID use, or sex, did not reveal any group differences in ratios of pT73-Rab10 to total Rab10 (Supplemental Fig. 5). These results suggest that PD cases with higher pT73-Rab10 to total Rab10 ratios have worse motor impairment, irrespective of disease duration.

### Elevated pT73-Rab10 to total Rab10 ratios in human serum are associated with antigenic pro-inflammatory responses in innate immune cells

Together with the serum samples, a second tube of whole blood was collected into PAXgene blood RNA tubes (IVD) for whole-transcriptomic analysis. Across the samples, *LRRK2* mRNA in whole blood is significantly but weakly correlated with LRRK2 protein levels in serum (r=0.119, p=0.007), but not correlated to the ratio of pT73-Rab10 to total Rab10 (r=0.008, p=0.848). To identify other genes and gene networks that might correlate with the ratio better, a weighted gene co-expression network analysis (WGCNA) identified two gene modules associated with the p773-Rab10 to total Rab10 ratio (Fig. 8A). The Purple gene module positively correlates with the ratio of pT73-Rab10 to total Rab10 as well as MDS-UPDRS III scores and LRRK2 levels in serum. The top-ranked cluster of interacting proteins encoded within the Purple module, identified by K-means clustering (mean local clustering coefficient 0.81, and Supplemental Fig. 6), highlights upregulated antigen presentation genes including *HLA-B*, *HLA-C*, and *HLA-E* (Fig. 8B). Variation in the *HLA-C* locus associates with PD-risk in genome-wide studies[47]. Differential gene expression analysis (DEG) demonstrates that mRNAs of several genes in the top-ranked Purple protein cluster (Fig. 8B) are upregulated in PD blood including *TYROBP* (log_2_FoldChange=0.22, Benjamini-Hochberg corrected p=5.07x10^-6^) and *CD14* (log_2_FoldChange=0.23, Benjamini-Hochberg corrected p=6.56x10^-5^). Other proteins in the Purple module cluster include CD14, a canonical surface marker for myeloid cells and coreceptor for toll-like receptors (TLRs), and the immunoreceptor tyrosine-based activation motif (ITAM), TYROBP, that regulates triggering receptor expressed on myeloid cells 2 protein (TREM2). In contrast to the Purple module, the ratio of pT73-Rab10 to total Rab10 is negatively correlated with the expression of genes in the Light Cyan module. The cluster of genes co-expressed in the Light Cyan module (mean local clustering coefficient = 0.808) highlights platelet integrin ITGA2B that connects to the platelet glycoprotein GP9, and interacts with P2RY12, a protein also associated with homeostasis in microglia. The Light Cyan cluster of genes are all highly expressed in platelets in blood and play central roles in immune interactions at the vascular wall (Fig. 8D). However, none of the Light Cyan cluster genes are differentially expressed between PD and controls in this cohort.

**Fig. 8.**
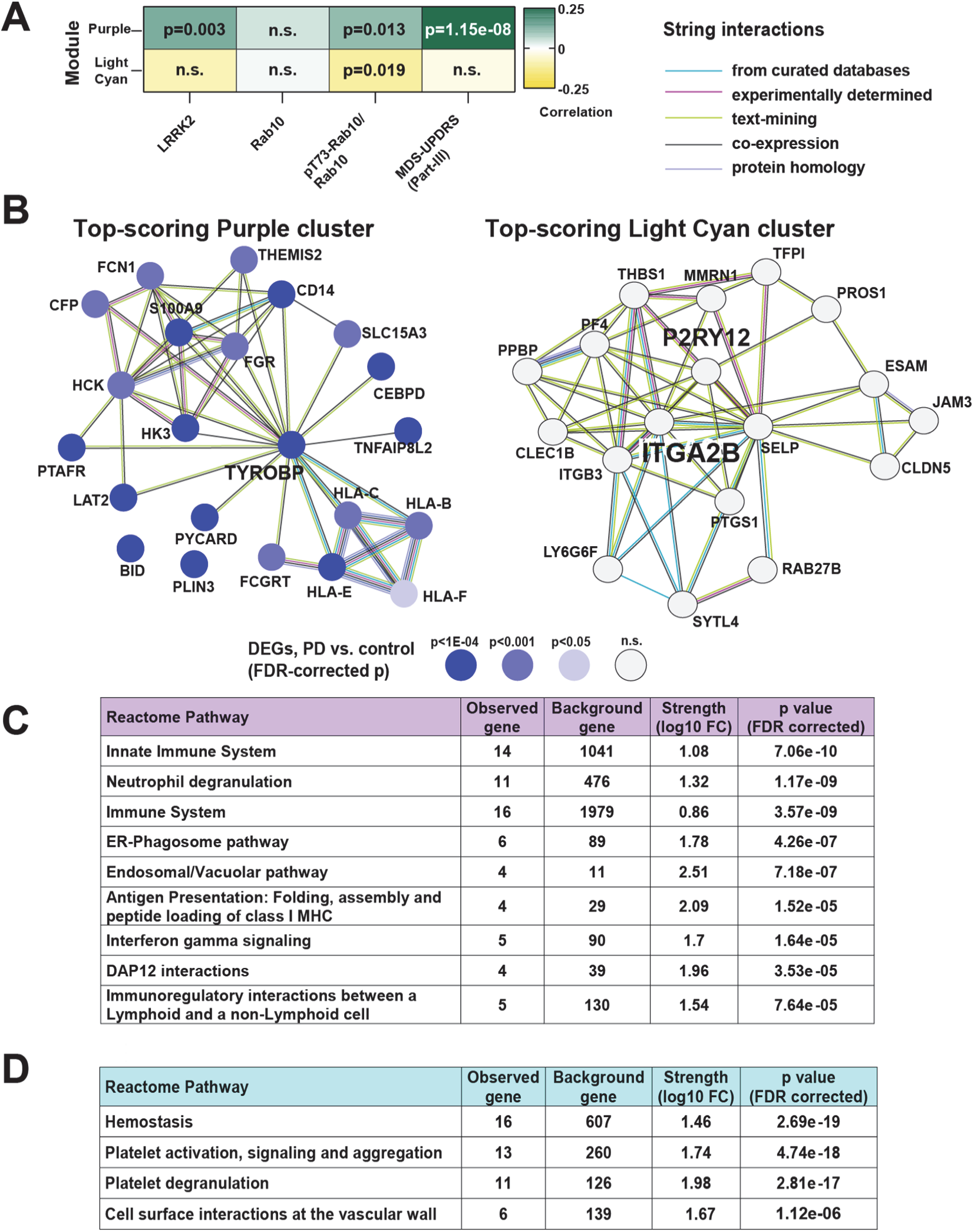
The ratio of pT73-Rab10 to total Rab10 in serum, and MDS-UPDRS III scores, are associated with myeloid cell activation and antigenic responses. **(A)** A weighted gene co-expression network analysis (WGCNA) identifies 27 gene modules of densely co-expressed genes (see Supplemental Fig. 6 for hierarchical clustering). Two modules, Purple and Light Cyan, are significantly correlated with the ratio of pT73-Rab10 to total Rab10. Pearson’s correlation coefficients and associated p values are additionally shown for correlation to LRRK2 protein levels in serum and MDS-UPDRS-III scores. **(B)** Top-scoring StringDB protein-protein interaction clusters from Purple and Light Cyan modules are shown. Differentially expressed genes (DEGs) in PD subjects vs. control are indicated from DESeq2 negative binomial fit, with level of significance (Benjamini-Hochberg adjusted p values) shown with a blue gradient. **(C)** All terms from the Reactome Pathway database (FDR-corrected p-value cutoff 1x10^-4^) are listed. Top process terms from Purple (N=290 searchable genes) and **(D)** Light Cyan (N=102 searchable genes) are shown. All terms with a p<0.05 cutoff from multiple databases are listed in Additional File 2.

## Discussion

The relationship between LRRK2-mediated Rab phosphorylation and disease progression has been poorly understood, in part due to the lack of sensitive and specific assays to evaluate these proteins that are low in concentration in most cells and biofluids. In different models, the LRRK2 substrate Rab10 has been implicated in immune functions related to both infection and PD[7,9,10,12,21]. As LRRK2 interacts with Rab proteins on vesicles, and some of these vesicles shed into the extracellular space, we utilized recent developments in single-molecule assay technology to design ultrasensitive assays to measure extracellular ratios of pT73-Rab10 to total Rab10. This is the first study we are aware of to evaluate hundreds of cases and controls for Rab phosphorylation, and a number of observations could be made in both rodent models and human samples. The ratio of pT73-Rab10 to total Rab10, as measured in biobanked serum samples, is highly dynamic and can rapidly diminish with oral administration of a LRRK2 inhibitor. These results may be compatible with evidence that extracellular vesicles and other components in serum shed from immune cells in blood may have very short half lives, for example ∼seven minutes in mice for exosome proteins[48]. In past studies in mice and rats, measures of LRRK2 kinase activity have typically required sacrificing the animal for western blot assessments of tissues, or procurement of cells from large volumes of blood. Since measurements from all three assays, total Rab10 and LRRK2, and pT73-Rab10, are accomplished with less than ∼60 microliters of serum, routine longitudinal measures in both clinical and rodent model samples are now possible.

Specificity of the assays is incredibly important in understanding the underlying biology of LRRK2 activity changes in different disease states and progression. The observed strong correlation between extracellular LRRK2 protein levels and pT73-Rab10 protein levels in serum across the cohort (r=0.70, p<0.0001) implies that most phosphorylated Rab10 in human serum is LRRK2-activity related. Evaluation of serum from clinical trial samples of LRRK2 kinase inhibitors will be informative to determine if all pT73-Rab10 signal from human sera is LRRK2-activity dependent, as it appears to be in the mouse models used in this study. Previous studies with clinical samples from LRRK2 inhibitor trials did not incorporate measures of the ratio of pT73-Rab10 to total Rab10[49,50], so it is difficult at present to compare our results in serum to other studies using different sources of protein. Following, a caveat to the interpretation of past studies is that the lack of total Rab10 measures to normalize the pT73-Rab10 levels may confuse lower total Rab10 levels with a reduction in LRRK2 kinase activity. For example, a diminished level of total Rab10 in serum, such as through immune cell depletion or that we observed in sepsis, may not necessarily reflect diminished LRRK2 activity should the ratio of pT73-Rab10 to total Rab10 persist at normal levels. In the human cohort of samples, total Rab10 protein levels positively correlate with pT73-Rab10 levels (in PD cases and controls, r=0.61, p<0.0001). Therefore, higher total Rab10 levels may erroneously drive the appearance of higher LRRK2 activity if only pT73-Rab10 is measured. Our data suggest prioritization of the ratio of pT73-Rab10 to total Rab10 as a truer metric of LRRK2 activity, since LRRK2 expression or activity does not appear to influence total Rab10 levels. Notably, the associated protocols used here are made available online via Protocols.Io in graphical step-by-step formats (see Methods), with commercially available parts and reagents. We expect the foundation results reported here together with the new assays will lead to a better understanding of how LRRK2 activity-dependent changes in the periphery may wax and wane in different disease models, and correlate with phenotypes in the brain. Samples already biobanked as part of ongoing clinical trials can also be rapidly assessed, as we have not observed significant freezer storage time effects on pT73-Rab10 levels.

Another important result from this study is that iPD patients with a very high ratio of pT73-Rab10 to total Rab10 in serum demonstrate worse clinical scores on the MDS-UPDRS III scale, as well as blood transcriptomic profiles that suggest elevated neutrophil and antigenic responses, compared to other iPD cases. The ratio of pT73-Rab10 to total Rab10 significantly predicts the expression of multiple *HLA* genes, including *HLA-C* that harbors variants associated with iPD risk[47]. Whether LRRK2 kinase activity and Rab phosphorylation regulates HLA traffic and antigen presentation in myeloid cells, or is responsive to these changes in immune cell maturation, is an area of ongoing investigation[51]. Though it is unclear whether LRRK2 activity itself is driving peripheral innate-immune cell inflammation, or responsive to it, there were no anti-inflammatory medications used by the participants in this study that appeared effective in reducing pT73-Rab10 levels. The reason for very high pT73-Rab10 levels found in some iPD patients is not known, and will be difficult to definitively pinpoint without intervention. In a recent study, we identified α-synuclein aggregates as a potent stimulator of pT73-Rab10 in monocytes, microglia and macrophages[12]. A critical clue in the mouse models studied here could be the high levels of pT73-Rab10 recorded in the serum of mice expressing wild-type *SNCA* at physiologically relevant levels[12,42,43]. We hypothesize that the high pT73-Rab10 signatures in *LRRK2-*negative iPD cases may be due to extensive and progressive peripheral and central α-synucleinopathy. Combining α-synuclein seeding assays in peripheral samples, serum, and CSF with LRRK2 activity biomarkers like pT73-Rab10 may help identify possible associations between these disease-associated features.

G2019S-*LRRK2* mutation carriers are known to have a more benign progressive motor phenotype over the course of disease[2]. Here, *LRRK2*-negative iPD carriers that show pT73-Rab10 levels in serum just as high (or above) that found in G2019S-*LRRK2* carriers have worse motor scores than other iPD cases with lower pT73-Rab10 levels. There are several potential explanations for these seemingly paradoxical observations. It can be presumed that the cause of upregulated pT73-Rab10 in *LRRK2-*negative iPD is fundamentally different from the cause of high pT73-Rab10 in G2019S-*LRRK2* carriers. The difference is exemplified in the mouse studies here showing that some septic wild-type mice developed similar pT73-Rab10 levels (or higher) than transgenic G2019S-*LRRK2* mice, and most wild-type septic mice showed higher serum pT73-Rab10 levels than R1441C-*Lrrk2* knockin mice. Though we did not perform behavioral evaluations of mice in this study, the septic mice are clearly sicker than transgenic G2019S-*LRRK2* mice, as expected. Elevated pT73-Rab10 in *LRRK2-*negative iPD may reflect a different but nevertheless detrimental disease process than that occurring in G2019S-*LRRK2* carriers. For example, recent studies show that many G2019S-*LRRK2* carriers are α-synuclein negative according to seeded-aggregation assays, with rarified α-synuclein pathology in the brain[52,53]. Since α-synuclein aggregates can drive pT73-Rab10 levels in immune cells[12], the difference in α-synuclein pathological progression between *LRRK2* mutation carriers and iPD may impact disease-related endpoints, even if pT73-Rab10 levels are comparable.

Fluid biomarkers in neurodegenerative diseases continue to have a transformative effect on understanding disease progression. Assays for pathological proteins neurofilament light chain (NFL[54]), phosphorylated tau (e.g., pTau-181[55]), and α-synuclein (both exosome-based[56] and seeding[57]), are now widely utilized in pre-clinical and clinical research. In contrast, much less is known about the biochemical activity associated with risk factors in iPD that include LRRK2 and glucocerebrosidase. While measures of phosphorylated Rab proteins or glucocerebrosidase activity are unlikely to be diagnostic for PD, they may provide critical information on how these activities change in disease, what drives these changes, and how best to therapeutically intervene.

## Conclusions

These data identify elevated LRRK2 kinase activity in the periphery, reflected in accurate measures of the ratio of pT73-Rab10 to total Rab10, as a biomarker for exacerbated inflammation. The type of inflammation associated with pT73-Rab10 to total Rab10 ratios, antigenic neutrophil responses, correlates with worsened disease endpoints in iPD. The assays provided here, ready for deployment in many laboratories and clinical sites across neurodegeneration research, will facilitate the discovery and advancement of new and effective LRRK2-targeting treatments to potentially help identify the patients most likely to benefit.

## List of abbreviations

PD: Parkinson’s disease
LRRK2: Leucine-rich Repeat Kinase 2
VPS35: vacuolar protein sorting 35
PBMC: peripheral-blood mononuclear cells
NFL: neurofilament light chain
SiMOA: single molecule assay
SDS: sodium deoxycholate
SNCA: synuclein-alpha
IFNγ: interferon gamma
TNF: tumor necrosis factor
PDBP: Parkinson’s Disease Biomarker Program
AMP-PD: The Accelerating Medicines Partnership Parkinson’s disease programme
DMR: Data Management Resource
MDS-UPDRS: Movement Disorders Society-Unified Parkinson’s Disease Rating Scale part I-IV;
MoCA: The Montreal Cognitive Assessment
UPSIT: University of Pennsylvania Smell Identification Test
LEDD: L-dopa Equivalent Daily Dosage
SD: standard deviation
SEM: standard error of the mean
IQR: interquartile range
CI: confidence interval
LL: lower limit of confidence interval
UL: upper limit of confidence interval.
BMDM: bone marrow-derived macrophage

## Declarations

### Ethics approval and consent to participate

All study protocols were approved by local Institutional Review Boards at the University of Alabama at Birmingham and Duke University, and Institutional Animal Care and Use Committees at Duke University and the Van Andel Institute.

### Consent for publication

N/A

### Availability of data and materials

Raw data are available for download online in Additional File 3.

### Author’s contributions

Y.Y, H.L., K.S., T.M., A.C., N.B., S.S, M.E., and R.B. performed experiments, analyzed data, and wrote the manuscript. L.H.S., M.W.L., D. V., and A.B.W. analyzed data and wrote the manuscript. T.A.Y., D.G.S., and D.J.M., contributed critical tools and wrote the manuscript. All authors read and approved the final manuscript.

## Acknowledgements

N/A

## Supplemental Figures

**Supplemental Fig. 1.**
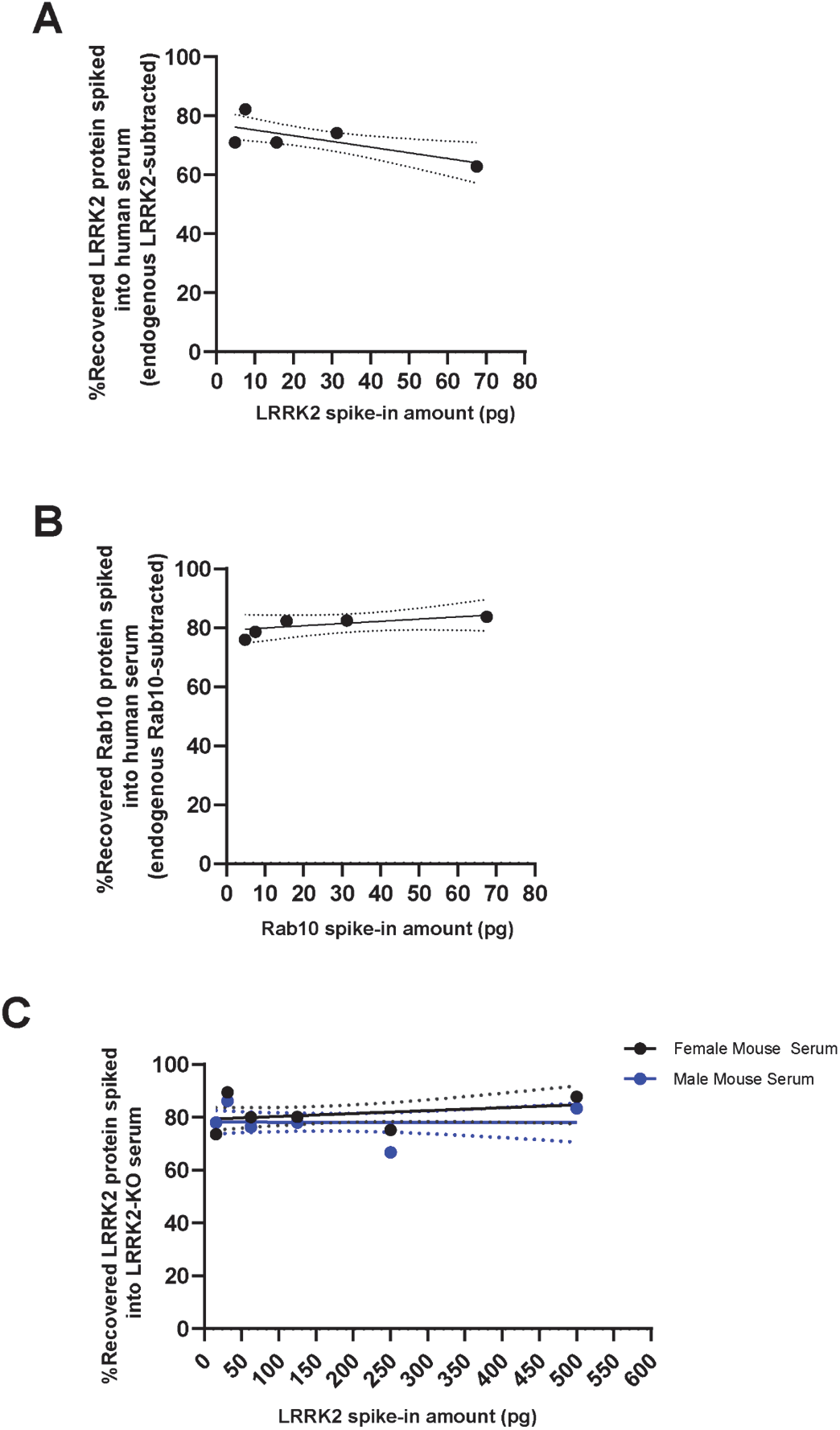
Spike and recovery results from recombinant proteins for LRRK2 and total Rab10 spikes into serum. **(A)** Lineplots show the percent recovery of picogram quantities of recombinant LRRK2 protein or **(B)** Rab10 protein spiked into human serum samples, with the background levels of endogenous LRRK2 or Rab10 subtracted prior to the calculated recovery. Mean recovery across the indicated range was 72.2%±9% for LRRK2 and 81.8%±7% for Rab10. **(C)** Recovery of recombinant LRRK2 protein in male (blue) or female (red) sera from LRRK2 knockout (LRRK2^-/-^) mice shows a similar recovery compared to human serum samples. Solid lines show mean linear regressions and dashed lines show 95% confidence intervals. Slopes in all plots are not significant according to correlation analysis.

**Supplemental Fig. 2.**
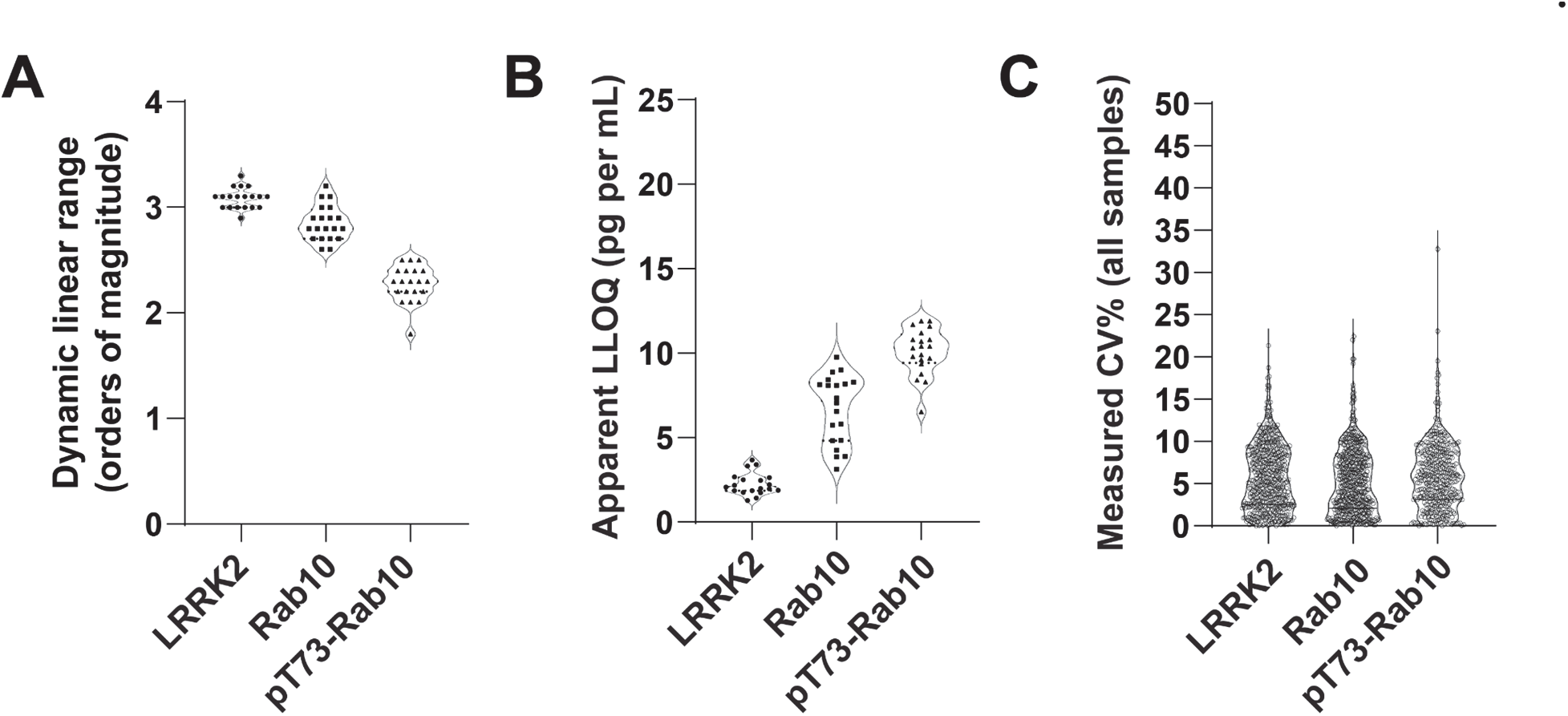
Good biomarker assay characteristics and reliability in biobanked human serum samples. **(A)** Column graph shows measured assay dynamic range calculated from at least nineteen different runs per analyte (i.e., LRRK2, Rab10, pT73-Rab10) on different plates. Mean observed dynamic range according to standard curves utilized in each run to resolve protein concentrations was consistent between runs and (in orders of magnitude) was calculated as 3.084 ± 0.095 S.D. for total LRRK2, 2.85 ± .16 S.D. for total Rab10, and 2.27 ± .17 S.D for pT73-Rab10. **(B)** Column graph highlights good assay reproducibility with respect to lower limits of quantification (LLOQ) across different runs. Each dot represents the LLOQ on a different plate from a unique run. The empirically determined lower limit of quantification (in pg per mL) for each run was consistent between runs and calculated as 2.257 ± 0.65 pg/mL S.D. for total LRRK2, 6.709 ± 2.01 pg/mL S.D. for total Rab10, and 10.10 ± 1.37 pg/mL S.D for pT73-Rab10. **(C)** Column graph provides evidence for excellent assay variance (CV, or coefficients of variability) across duplicated or triplicated human serum samples. Each dot shows the calculated CV calculated from one human serum sample. The average (mean) CVs are 5.816 ± 3.86 S.D. for measurements of total LRRK2, 5.361 ± 3.9 S.D. for measurements of total Rab10, and 5.906 ± 3.95 S.D for CVs related to the measures of pT73-Rab10.

**Supplemental Fig. 3.**
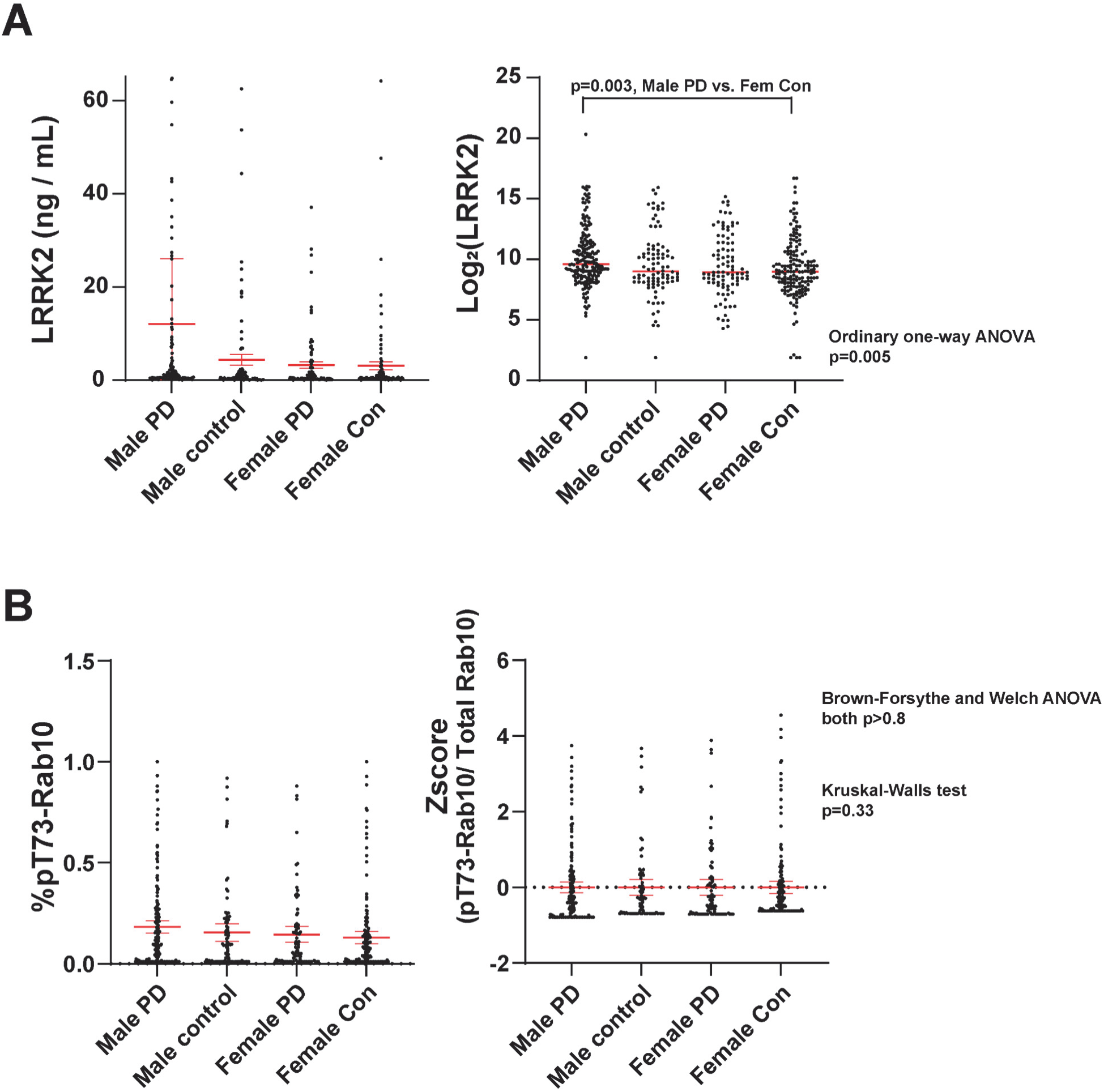
Serum LRRK2 and pT73-Rab10 to total Rab10 ratios are variable across subjects and similar in PD and control subjects. **(A)** Scatter column plots show measured (unadjusted) LRRK2 levels in serum from male and female PD and control, and Log(2) transformation results that result in a normal distribution of LRRK2 expression in the groups. An ordinary one-way ANOVA is significant, with Tukey’s post-hoc test demonstrating nominally increased levels of LRRK2 in male PD subjects versus control female subjects. **(B)** Scatter column plots show measured (unadjusted) ratios of pT73-Rab10 to total Rab10 in serum from male and female PD and control. Attempts to transform the data into a normal distribution were not successful, and shown is a Z-score transformation that highlights the strata of subjects with very low (<0.09%) Rab10 phosphorylation (i.e., the ratio of pT73-Rab10 to total Rab10), in addition to some subjects with very high pT73-Rab10 levels. ANOVA test results do not support an overall difference in pT73-Rab10 ratios between groups. Together, these results suggest that LRRK2 and pT73-Rab10 levels in serum are unlikely to be useful for diagnostic purposes in the separation of PD cases from controls.

**Supplemental Fig. 4.**
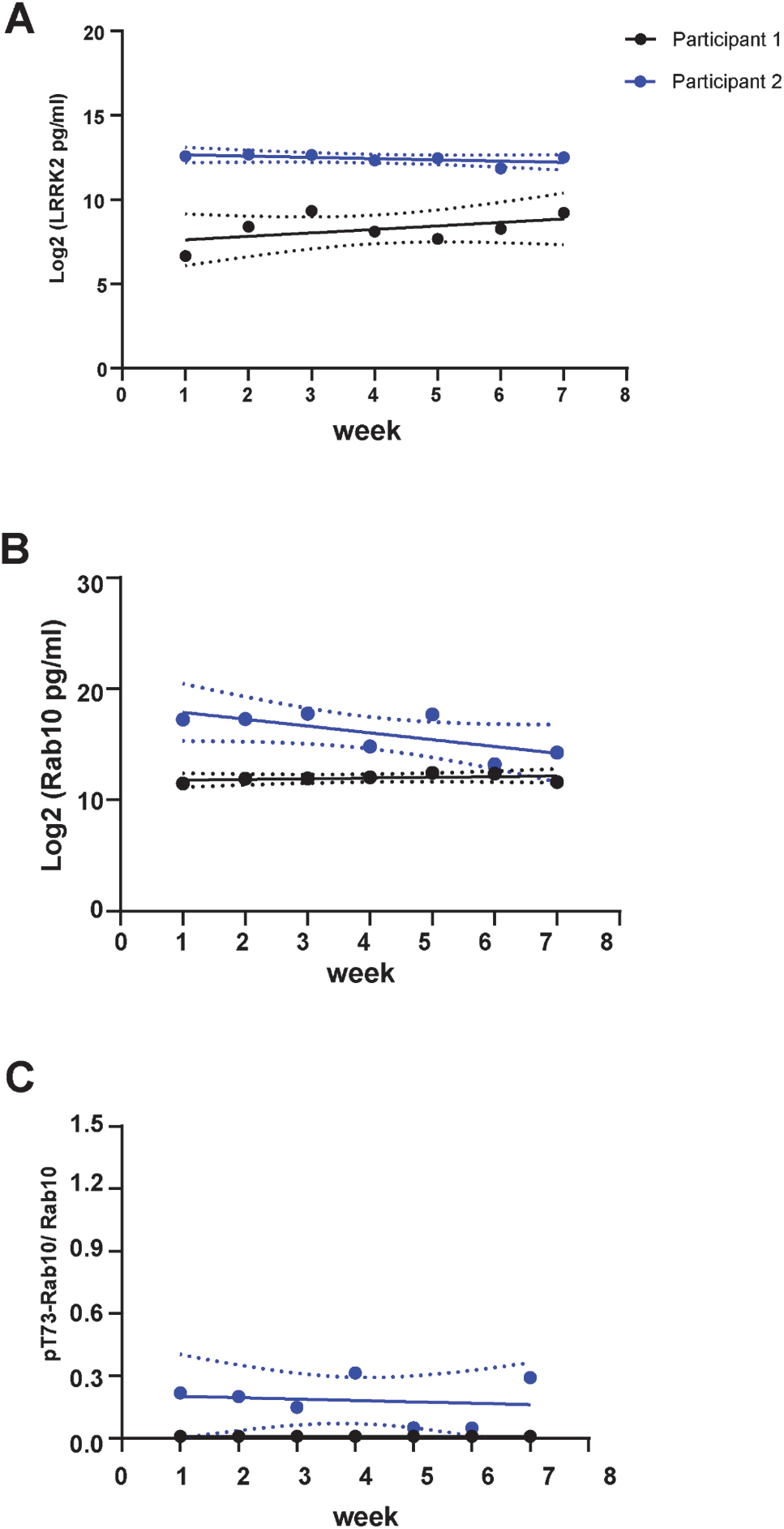
Longitudinal data (∼2 months) for serum biomarkers from weekly serum depositions from 2 healthy control participants recruited in the PDBP cohort. **(A)** Line graphs show LRRK2 expression sampling repeated with 1 week intervals for two control subjects recruited in the PDBP. Samples were run across different plates and randomized with final data curation prior to unblinding the samples (see Methods). **(B)** Total Rab10 levels over time, and **(C)** the ratio of pT73-Rab10 to total Rab10 over time. Solid lines show mean linear regressions and dashed lines show 95% confidence intervals. Slopes in all plots are not significant according to correlation analysis.

**Supplemental Fig. 5.**
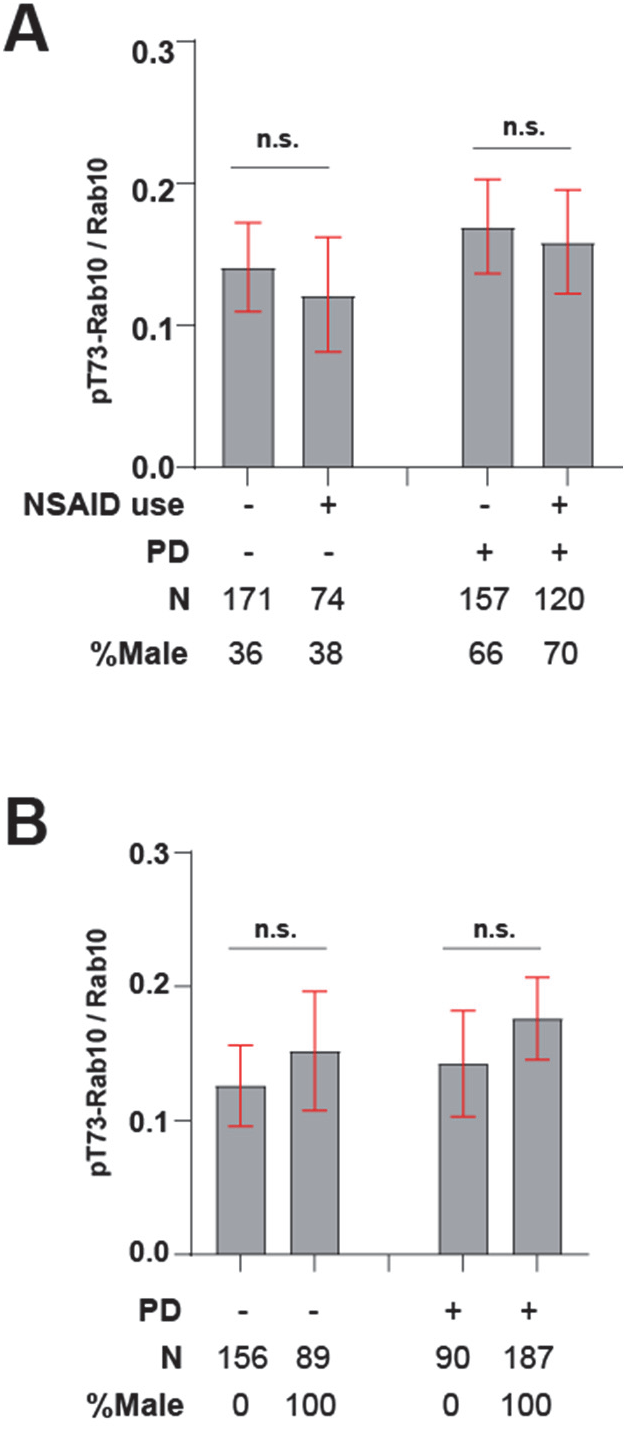
No interactions between the ratio of pT73-Rab10 to total Rab10 in NSAID use or sex as covariables. Serum levels of total LRRK2 and pT73-Rab10/Rab10 ratios binned according to **(A)** usage of non-steroidal anti-inflammatory medication or **(B)** PD diagnosis and sex. The proportion of participants in each group diagnosed with PD is indicated. Column graphs show group means with 95% confidence intervals (C.I.) as red-colored error bars. p values are from Mann-Whitney tests for comparisons of the unequal group sizes, and n.s. is not significant (p>0.05).

**Supplemental Fig. 6.**
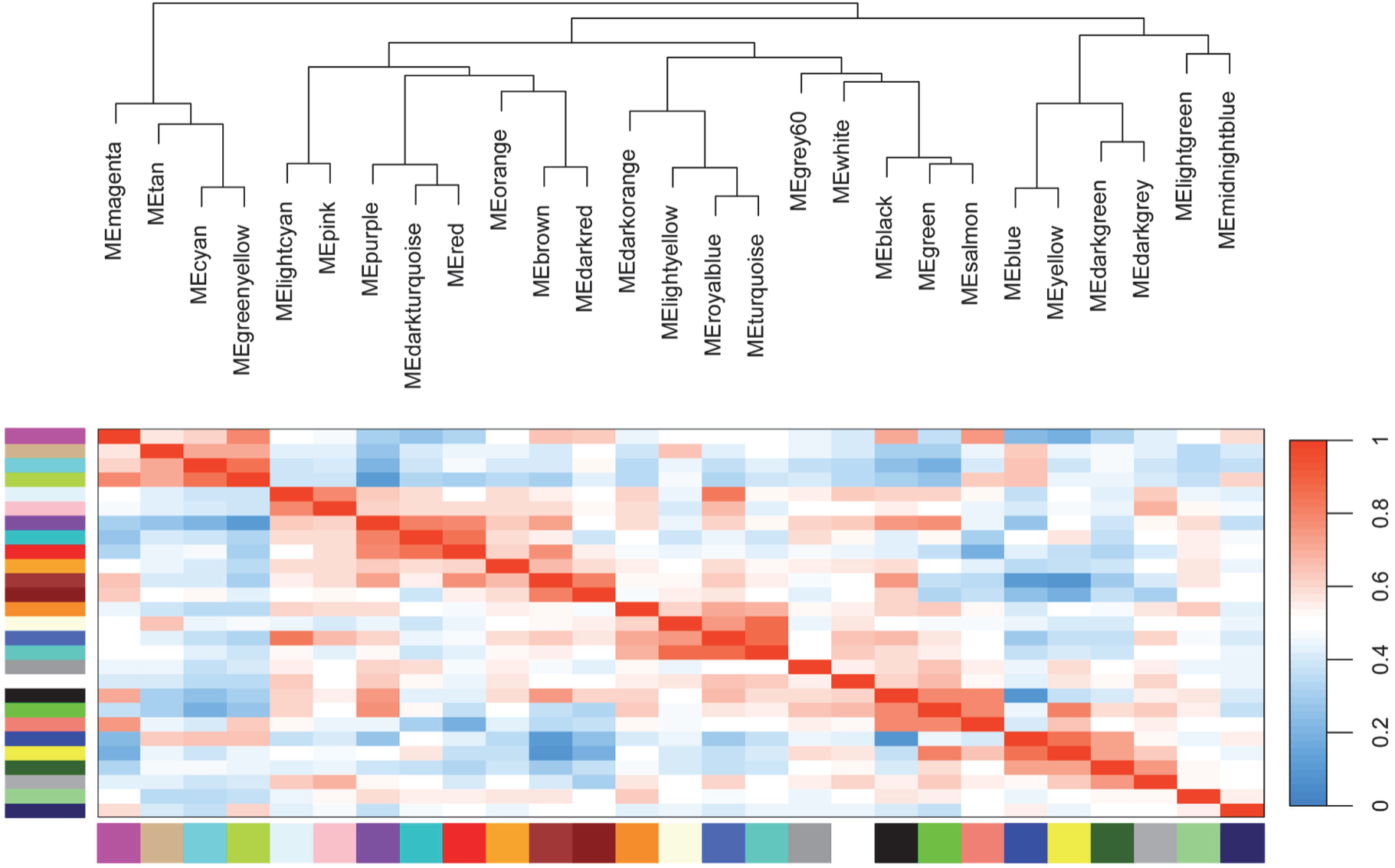
Hierarchical clustering and adjacency between modules identified in weighted correlation network analysis (WGCNA) of PD cases and controls (Table I). The R library “WGCNA” was utilized to visualize modules of highly correlated genes from variance-stabilizing transformed transcriptomic data in whole blood samples. A total of 27 gene modules were produced. In order to study the relationships among the found modules, the eigengenes, or the first principal component of each module, were used to represent the gene expression profile and module similarity was quantified by correlating the eigengene values. The *plotEigengeneNetwork* function was used to generate a dendrogram and a corresponding heatmap. Hierarchical clustering of the eigengenes visualizes similarity. The modules of interest with respect to pT73-Rab10 to total Rab10 ratios, Purple and Light Cyan, are distantly related. The heatmap visualizes eigengene adjacency (A_IJ_ = (1+cor(*E_I_, E_J_*))/2), with higher values (red shades) indicating greater similarity. Purple and Light Cyan modules are similar to a moderate extent (adjacency = 0.63).

